# Massively parallel characterization of RNA G-quadruplex stability and molecular recognition

**DOI:** 10.1101/2025.04.29.651304

**Authors:** Justin G. Martyr, Bryan B. Guzmán, Yue Hu, Rebekah L. Rothacher, Anthony M. Mustoe, Maria M. Aleman, Daniel Dominguez

## Abstract

RNA G-quadruplexes (rG4s) have been implicated as important regulators of RNA metabolism and are promising targets for RNA-targeted therapeutics. rG4s typically require a canonical (G_≥2_N_1-7_)_4_ motif, but the sequence features that affect rG4 stability and recognition by RNA-binding proteins (RBPs) and rG4-binding ligands are not fully understood. To interrogate sequence-level drivers of rG4 folding, we applied a reverse-transcriptase stop sequencing strategy to a library of ∼3,000 synthetic rG4s with varied G-tract lengths, loop lengths, and loop compositions, permitting massively parallel quantification of rG4 stability. Our data confirm known sequence-level features and characterize novel combinatorial impacts of these features. We also assessed systematically mutagenized natural rG4s, revealing unexpected mutations that significantly affect rG4 stability, including contributions from flanking sequences outside of the rG4. We further used our strategy to assess rG4 recognition preferences of the model rG4- ligand pyridostatin, revealing a preferential stabilization of rG4s containing mixed-length G-tracts. We further demonstrated the potential for large-scale protein binding assays with our library to reveal rG4 features recognized by RBPs, specifically G3BP1 and FMRP. Our approach and data provide a generalizable framework to study sequence-level drivers of rG4 stability, binding by RBPs, and ligand interactions, defining basic principles of rG4 formation and downstream biology.

## INTRODUCTION

Regulation of RNA processing and downstream gene expression has increasingly been linked to structured RNA elements (1,2). These elements, in part through their interactions with RNA binding proteins (RBPs), impact transcription (3,4), as well as co-transcriptional (5,6) and post-transcriptional RNA processing (7,8). Thus, understanding and predicting sequence features that drive RNA structure formation is of high importance but remains a significant challenge.

RNA G-quadruplexes (rG4s) are regulatory structured elements that arise from guanine-rich sequences (9–11) and are over-represented in the transcriptome (12,13). rG4s form from guanines arranged in planar, Hoogsteen-bonded G-tetrads that stack on each other and coordinate a metal ion, typically potassium (**Fig. 1A**) (Reviewed in: (14)). rG4 folds are highly stable (10), a feature likely dependent on their sequence composition. The underlying sequence motif for rG4 forming sequences is generally defined as (G_≥2_N_1-7_)_4_, yet this pattern allows for an extraordinary level of diversity (15). Despite some understanding of general rG4 trends (16–18), this massive diversity makes the relationship between rG4 sequence and stability challenging to study. Moreover, sequence contexts outside of rG4 forming sequences and their impact on folding remain largely unknown. A nuanced evaluation of rG4 stability and recognition is needed to understand how sequence variants (*e.g.*, polymorphisms) impact these structures and to build predictive models for these sequence-level drivers.

**Figure 1.**
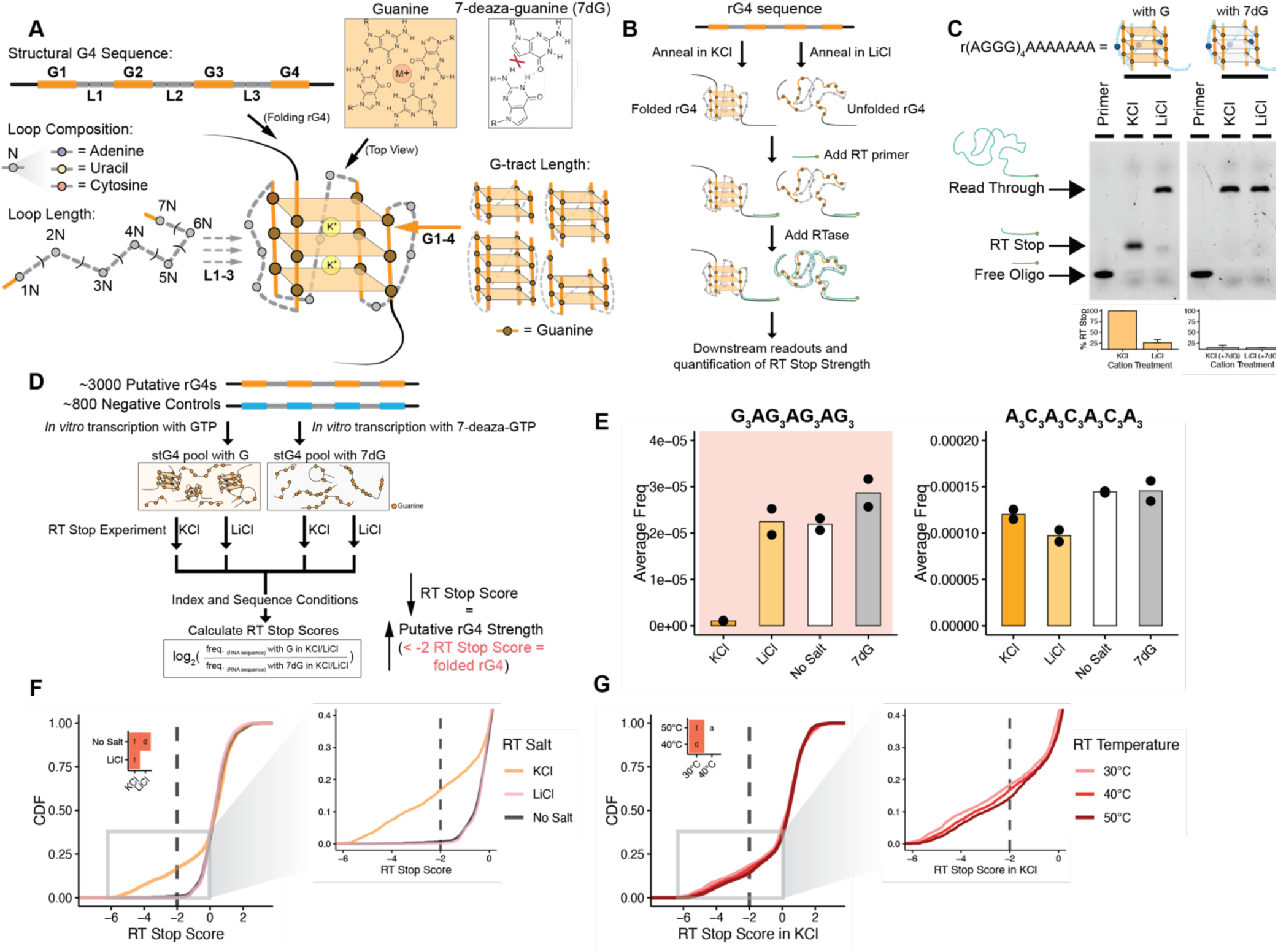
Design and initial characterization of Structural rG4 Pool. **A)** Schematic of RNA G-quadruplex sequence features. The design of our structural rG4 pool focused on using the base rG4 sequence and varying the G-tract lengths, loop nucleotide composition, and loop length. This pool was *in vitro* transcribed with either guanine or 7-deaza-guanine (7dG), which is unable to fold into an rG4. **B)** Schematic of RT stop experimental workflow. **C)** Evaluation of RT stop protocol against a strong rG4 (r(AGGG)_4_AAAAAAA) either prepared with guanine or 7dG. Representative gel for RT stop products and full-length read through products and %RT Stop was quantified (n=2). **D)** Schematic of RT stop sequencing procedure and metric for evaluating RT Stop Scores. **E)** Bar graphs of a putative rG4 (G_3_AG_3_AG_3_AG_3_, red background) and a negative control (A_3_C_3_A_3_C_3_A_3_C_3_A_3_) and their respective average frequencies in KCl (orange), LiCl (light orange), No Salt (white) and with 7dG (grey) (n=2). Cumulative distribution function (CDF) of **F)** RT Stop Scores at 40°C with different salt conditions (KCl in orange, LiCl in pink, and No Salt in black) and **G)** RT Stop Scores in KCl with different RT temperature conditions (30°C in light red, 40°C in red, and 50°C in dark red) (n=2). Plot insets (*left*) show p-values determined by two- sided KS test corrected by BH procedure. Red square indicates p≤0.05. Values are as follows: a (ns), d (p≤0.01), f (p≤0.0001). Zoomed in plots (*right*) shows RT Stop Scores less than 0.

rG4s have been found to influence several RNA processing and regulatory steps including localization (7), translation (19–21), and splicing (6,22). rG4s have further been implicated in stress granule formation (23), protein folding (24,25), and even linked to disease pathways, especially those involved in neurodegenerative diseases (26,27) and cancer (Reviewed in: (28)). The prevalence of rG4 folding in cells remains an open question, with one study indicating rG4s are unfolded in cells (29) and another study, using the same methodology, indicating rG4s fold in response to cellular stress (23).

The function of rG4s likely depends in part on their ability to interact with RBPs. Many rG4-binding RBPs have been identified (30–36), however, the methods used focus primarily on RBP interactions with select rG4 sequences, leaving RBP specificity for rG4s largely unclear. Mechanistic insight into how proteins bind rG4s is limited, especially with respect to structural studies. This deficiency is likely because intrinsically disordered domains appear to be primary drivers of rG4 binding (Reviewed in: (37)), complicating structural assessment and definition of basic biochemical principles which drive these interactions.

The rG4 structure has also become a prominent target for small molecule ligands and nucleic acid therapeutics (38–40). Targeting rG4s on the transcriptomic level with both stabilizing and destabilizing ligands has led to the identification of hundreds of genes that are differentially expressed (41). Additionally, small molecule treatments have been found to have differential impacts on specific RBP-rG4 interactions (42). While the focus of the field has turned towards targeting specific disease-relevant rG4s (*e.g.*, SARS-CoV-2 (43), dystrophin in Duchene Muscular Dystrophy (44)), the characteristics that make rG4s ‘targetable’ or ‘druggable’ are still not well understood. Conversely, both structural and sequence features that are recognized by known rG4 ligands remain to be fully characterized.

The remarkable stability of some rG4s is known to inhibit the processivity of enzymes when folded. For example, rG4 formation can inhibit ribosomal readthrough (Reviewed in: (45)). Helicases can counteract this stability via directed unfolding and have demonstrated *in cellulo* activity in unfolding rG4s (46,47). rG4s can also inhibit reverse-transcriptases (RTases), which are often unable to reverse transcribe through rG4-containing sequences. This feature has birthed a suite of techniques to identify rG4 forming sequences across the transcriptome by identifying sequences that trigger an ‘RT stop’ (29,48–50). RT stop experiments have also been used to characterize small molecule and mutational impacts on rG4 folding (51–53).

To directly evaluate sequence-level drivers of rG4 folding, we designed a library of ∼3,000 synthetic rG4s with varied G-tract lengths, loop lengths, and loop compositions, as well as 6 natural sequence rG4s which we tiled with nucleotide mutations. By RT stop sequencing, we confirm known characteristics which increase rG4 stability and further reveal loop composition preferences and combinatorial variations of these features which can similarly modulate rG4 folding stability. In selected natural rG4s, we point to specific mutations that alone drastically shift that specific rG4’s stability, leading to novel insights into natural rG4 folding. We also assess rG4 recognition by the model rG4 ligand, pyridostatin (PDS), a perceived global G4 stabilizer (54,55), and identified small molecule-induced changes in stability driven by specific underlying sequences. Finally, through the use of a massively parallel assay, RNA Bind-n-Seq (56), we highlight specific rG4 variants bound by known rG4-binding RBPs, and decipher general rG4 characteristics which influence binding. Our approach enables derivation of principles for rG4 folding and improves our understanding of fundamental rG4 recognition and biology.

## RESULTS

### Design of an rG4 pool to systematically evaluate principles of rG4 folding stability

RNA G-quadruplexes (rG4s) are well-known to cause premature termination during reverse transcription (RT), a feature that has been exploited to identify rG4 structures at the transcriptomic level (48) and evaluate impacts of ligands on rG4 stability (42,53). As a proof of principle, we performed an RT stop experiment on a strong rG4 (r(AGGG)_4_) with a FAM-labelled RT-oligo (**Fig. 1B**). rG4 stability can be shifted using cations, either using K^+^ to stabilize or Li^+^ to destabilize rG4s (14). Resolving RT products by gel electrophoresis showed an RT stop in KCl, and a readthrough product when LiCl was present (**Fig. 1C**, *left*). We also *in vitro* transcribed the strong rG4 sequence with 7-deaza-guanine (7dG) instead of guanine. 7dG has been documented to prevent rG4 folding due to a lack of hydrogen bonding capabilities at the N7 position (57,58) (**Fig. 1A**). No RT stop product was detected for the 7dG strong rG4, confirming that 7dG rG4 sequences lack RT stop activity (**Fig. 1C**, *right*).

We sought to exploit these features to enable massively parallel measurement of rG4 stability by sequencing. We expected that folded rG4s would cause RT-termination proportional to the stability of the rG4, thus leading to lower reverse transcribed RNA. Therefore, specific RT conditions which shift rG4 stability would differentially result in sequence dropout after polymerase chain reaction (PCR). Sequencing the readthrough products and comparing across RT conditions would then allow for high throughput quantification of rG4 strength. We designed a 6,000 oligo RNA pool composed of rG4 and non-rG4 forming sequences. This pool includes a subpopulation of synthetic putative rG4s derived from the canonical rG4 sequence, (G_≥2_N_1-7_)_4_ (**Fig. 1A**). We included variations across three main components, specifically: **i**) length of G-tracts, **ii**) length of loop sequences, and **iii**) nucleotide composition of loops (**Fig. 1A**). We allowed G-tract length to vary between GG, GGG, GGGG, leading to the formation of two, three, and four tetrad rG4s, as well as G-tract combinations of different lengths. For loops, we selected polyA, polyC, and polyU, and varied the length between 1-7 nts. We also allowed different combinations of polyA and polyU loops with varying lengths (*i.e.*, GGG<u>AAA</u>GGG<u>UUU</u>GGG<u>A</u>GGG). In total, this accounted for 3,213 unique putative rG4 sequences (**Table S1**). This structural rG4 (stG4) pool additionally included a small series of polyUG (pUG) sequences that have been shown to adopt a G-quadruplex fold, and a set of non-rG4 controls including synthetic A-rich sequences and natural sequences with varied GC-content but not predicted to form rG4s. The pool also included select natural sequence rG4s derived from human mRNAs (to be discussed below) for a total of 6,000 sequences (**Methods**).

### Sequencing RT stop allows for massively parallel quantification of rG4 strength

We combined RT stop assays with high throughput sequencing using our designed stG4 pool (**Fig. 1D**). RT stop profiling using this pool has some key advantages: **i**) all RNAs were synthesized to be equal in length, **ii**) we can control RT reaction conditions including cation concentrations and temperatures, **iii**) all sequences were amplified using the same primer sets limiting potential RT and PCR differences related to priming, and **iv**) use of 7dG RNA pools provided a way to control for rG4 sequences by promoting RT readthrough through inhibition of rG4 folding (**Methods**). To assess RNA folding, we *in vitro* transcribed the stG4 pool with either guanine or 7dG and performed RT stop experiments in KCl or LiCl (**Fig. 1D**). After normalization for sequencing library size (typically ∼10 million reads), we calculated an ‘RT Stop Score’ by taking the log_2_ fold change of the frequency of a given RNA sequence in the guanine stG4 pool to the same sequence’s frequency from the 7dG stG4 pool (**Fig. 1D**). We expected strong rG4 sequence frequencies to be low in guanine-containing RNAs in KCl compared to 7dG-containing RNAs. As an initial quality check, we compared sequence frequencies across RNA replicates with guanine and 7dG in KCl. We found high positive correlations of individual sequence frequencies, and as expected folded rG4s had lower frequencies with guanine and KCl when compared to those made with 7dG (**Fig. S1A**, *left, middle*). This led to a strong correlation between RT Stop Scores, indicating this to be a reproducible assay (**Fig. S1A**, *right*).

To compare, we set a threshold of an RT Stop Score of -2 as a ‘Folded rG4’ cutoff, meaning the frequency of the given rG4 sequence made with guanine is 4-fold lower than the frequency of the same RNA with 7dG (**Fig. 1D**). G_3_AG_3_AG_3_AG_3_, the same rG4 motif highlighted in our proof of principle RT stop experiment (**Fig. 1C**), displayed markedly lower frequency in KCl compared to LiCl, a no salt condition (no salt added), or in RNA made with 7dG, confirming the expected result (**Fig. 1E**, *left*). Conversely, an example non-rG4 A_3_C_3_A_3_C_3_A_3_C_3_A_3_ showed no trend with minimal differences across RT conditions. (**Fig. 1E**, *right*). It should be noted that RT Stop Scores were on a spectrum and normalized, so scores from -2 to 0 (and potentially greater than 0) suggest weaker rG4 folding, but do not necessarily rule out dynamic or weak rG4s that were more readily unfolded by RTases (discussed in more detail below). Filtering our data using different read cutoffs (per RNA sequence) demonstrated excellent correlations and robustness across all cutoff criteria (10-200 reads per RNA) (**Fig. S1B**). However, significant sequence dropouts were observed in KCl libraries, an expected result due to strong rG4s triggering RT stops and yielding minimal product for sequencing. The strong rG4 drop-out was substantially reduced in LiCl or 7dG library conditions (**Fig. S1C**).

We next evaluated RT Stop Scores across multiple RT conditions, including the presence or absence of 100 mM monovalent cations (K^+^, Li^+^) during the RNA folding and RT steps, and the impact of different RT temperatures (30 °C, 40 °C, and 50 °C) (**Fig. 1F, G**). As expected, K^+^-buffer revealed a distribution of negative RT Stop Scores, consistent with K^+^-dependent rG4 folding. The majority of RNAs had weak, positive RT Stop Scores in LiCl or in no salt conditions, but a limited number of RNAs had RT Stop Scores less than 0, suggesting some folding of rG4 structures even in the absence of K^+^ (**Fig. 1F**, and discussed below). Increasing the RT extension temperature from 30 °C to 50 °C resulted in a modest but significant increase in RT Stop Scores, consistent with destabilization of strong rG4s at higher temperatures (**Fig. 1G, Fig. S1D, E**). Taken altogether, this demonstrates the robustness and reproducibility of our approach in characterizing rG4 strength in parallel with different RT conditions.

### G-tract length is a critical driver of rG4 strength

Lower-throughput biochemical assays have demonstrated that G-tract length and intervening loop length directly influence rG4 stability (16,17,59,60). As a first pass, these trends were well-captured in our analysis. Stratifying our pool by either putative rG4s or non-rG4 controls, we found a significant left-shift (more folding) in RT Stop Scores for putative rG4s in KCl (**Fig. 2A**). This effect was weaker in LiCl, consistent with K^+^-dependent rG4 stabilization (**Fig. S2A**). Stratifying putative rG4s by minimum G-tract length (*i.e.*, classifying each rG4 by its shortest G-tract: GG, GGG, or GGGG) revealed that GGGG-rG4s generated the strongest, most negative RT Stop Scores (**Fig. 2B**). Most GGG-rG4s also showed negative RT Stop Scores, but to a lesser extent than GGGG-rG4s. By comparison, the majority of evaluated GG-rG4s demonstrated positive RT Stop Scores, suggesting that they were less able to fold in these experimental conditions. This effect may also reflect RT Stop assay-specific signal; that is, GG-rG4s may fold but not strongly enough to inhibit RTases. Interestingly, while RT Stop Scores were substantially weakened in LiCl, we still observed G-tract length dependence, consistent with the strongest rG4s folding even in LiCl (**Fig. S2B**). As noted above, we included sequences containing mixed tract lengths (*e.g.*, GGGN<u>GG</u>NGGGNGGG, 3-2-3-3) enabling interrogation of more complex rG4 forming sequences. Comparison of maximum G-tract length rather than minimum G-tract length revealed that the stability of an rG4 was largely determined by the shortest G-tract within the rG4 (**Fig. 2C**).

**Figure 2.**
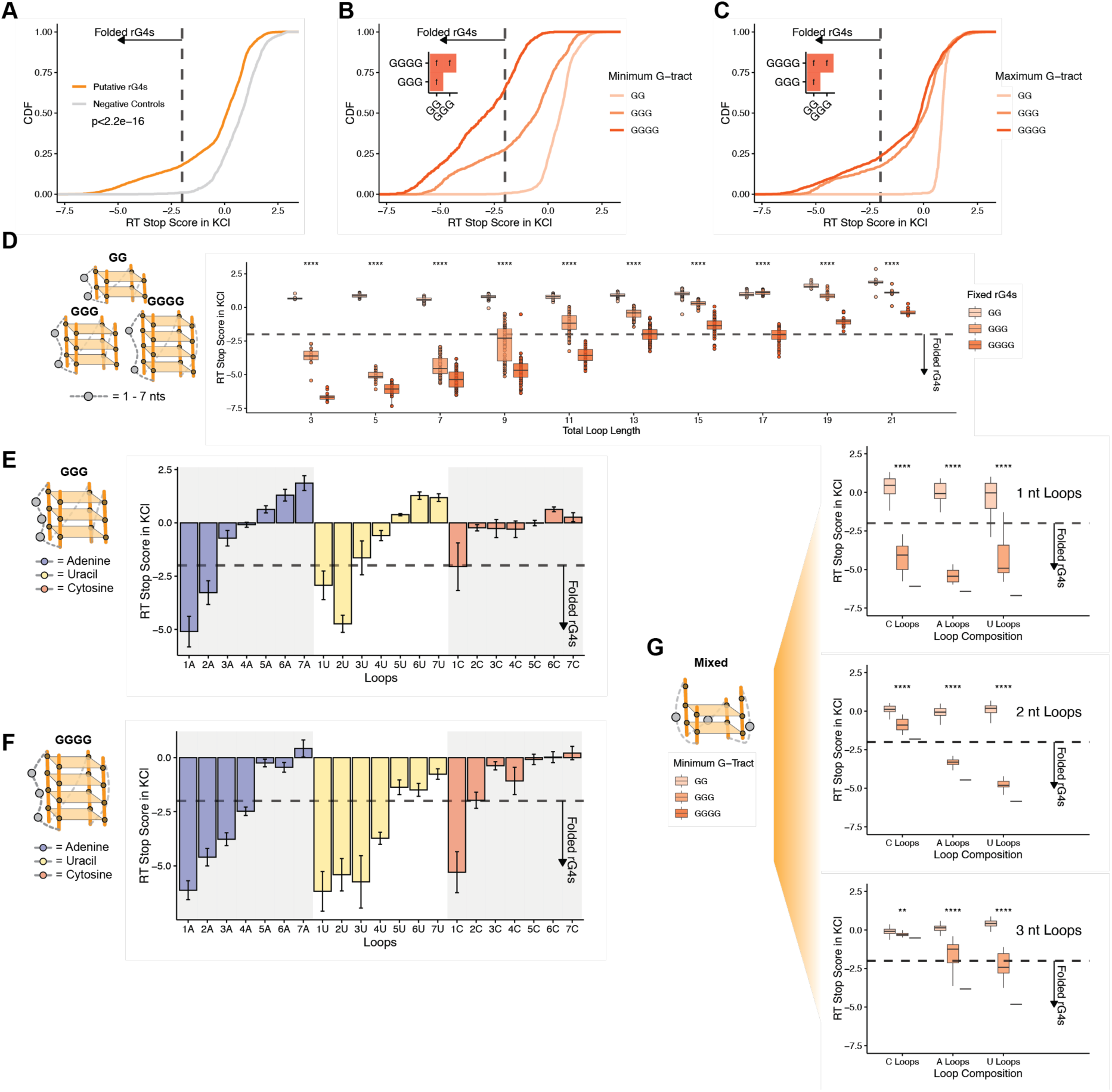
Evaluating characteristics of rG4 folding. CDF of the structural rG4 pool oligo RT Stop Score in KCl **A)** by putative rG4s (orange), and negative controls (grey) (Data is the average of 3 independent replicates), then further **B)** by minimum and **C)** by maximum G-tract length, specifically GG (light orange), GGG (orange), and GGGG (dark orange) (Data is the average of 3 independent replicates). Line at -2 RT Stop Score is indicative of cutoff for folded rG4s. P-value is shown, or plot insets (*left*) show p-values determined by two-sided KS test corrected by BH procedure. Red square indicates p≤0.05. Values are as follows: f (p≤0.0001). **D)** G-quadruplex schematic (*left*) demonstrating selection of oligos locked for GG, GGG, and GGGG at all G-tract positions (fixed rG4s) and varying the loop length. Box plot of RT Stop Score in KCl (*right*) for oligos having fixed G-tract length of GG (light orange), GGG (orange), GGGG (dark orange) with the total loop length (sum of nucleotides across all loop lengths) (Data is the average of 3 independent replicates). Line at -2 RT Stop Score is indicative of cutoff for folded rG4s. P-values show significance of the distribution of means by minimum G-tract, **** = p≤0.0001. Groups with less than 8 sequences were not included in analysis. Bar plots for only **E)** GGG and **F)** GGGG putative rG4s and the RT Stop Scores in KCl with differing loop lengths and compositions. This is 1 nt to 7 nt loops of the same nucleotide composition (i.e., 1A, 2A, 3A…) with A loops in blue, U loops in yellow, and C loops in red. Line at -2 RT Stop Score is indicative of cutoff for folded rG4s. **G)** Box plots for distribution of RT Stop Score in KCl for all G-tract permutations (total 81 unique combinations: 65 GG, 15 GGG, 1 GGGG) per loop composition with either 1 nt (*top*), 2 nt (*middle*), and 3 nt (*bottom*) loops. Colors show minimum G-tract length, specifically GG (light orange), GGG (orange), GGGG (dark orange). Line at -2 RT Stop Score is indicative of cutoff for folded rG4s. P-values show significance of the distribution of means by minimum G-tract: ** = p≤0.01, **** = p≤0.0001.

### Loop analysis reveals length and composition biases for rG4 folding

As an initial evaluation of loop contributions, we first totaled loop length (the sum of all loops within an rG4) across only fixed G-tract rG4s (*i.e.*, all tracts were GG, GGG, or GGGG) to correlate loop length and G-tracts combinatorially. Analysis of total loop length showed the strongest stability for 1 nt loops (3 total nts) (**Fig. 2D**). Additionally, rG4s with longer G-tracts were better able to tolerate longer loops (**Fig. 2D**), but loops greater than 13 total nts were able to disrupt the folding of even most GGGG-rG4s. To our knowledge, this is the first comprehensive assessment evaluating the interplay between G-tract length and loop length for rG4 stability.

We next assessed the nucleotide composition of loops to define novel insights into rG4 stability. A and U loops had variable impacts on rG4 folding whereas C loops largely led to unfolding (**Fig. S2D**). For a more nuanced understanding of how loop composition interplays with findings described above, we compared RT Stop Scores across fixed G-tract rG4s (GGG: **Fig. 2E**, GGGG: **Fig. 2F**) with increasing loop lengths of either A/U/C nucleotide compositions. GGG- and GGGG-rG4s demonstrated that increasing A loop length led to consistent rG4 destabilization (**Fig. 2E, F**). Increasing U loop length had a lesser destabilizing effect; in fact, increasing U loop length from 1 nt to 2 nt enhanced stability for GGG rG4s (**Fig. 2E, F**). rG4s composed of only GG G-tracts (**Fig. S2E**) and C loops longer than 1 nt demonstrated positive RT Stop Scores, therefore weakly folding.

We next evaluated short and U loop stabilization in the context of mixed G-tract rG4s (*i.e.*, containing combinations of GG, GGG, and GGGG G-tracts) (**Fig. 2G**). While 1 nt loops had similar strengths for each minimum G-tract, increasing loop lengths recapitulated the findings above as U loops were generally stronger than the corresponding A and C loops even in mixed rG4s (**Fig. 2G**, *middle, bottom*). Altogether, these data highlight the combinatorial tradeoffs that govern rG4 stability.

### Mutating natural sequence rG4s provides nucleotide-level insights into rG4 folding

Unlike the simplified model rG4s designed above, natural rG4 sequence compositions are complex in and outside of the rG4 forming region. Furthermore, many RNA regions contain multiple potential rG4 forming G-tracts, which make defining folding topology difficult using available approaches. To further delve into sequence-level drivers of natural rG4 folding, we selected six natural sequence rG4s with varied predicted propensities for rG4 folding using RNAFold (61). Specifically, three strong rG4s (*VEGFA*, *U2AF2*, *ARHGEF12*), two weaker rG4s (*EWSR1*, *HNRNPA2B1*), and a non-rG4 (*PABPN1*) were chosen (**Fig. 3A**). As expected, wild-type (WT) sequences for strong rG4s *VEGFA*, *U2AF2*, and *ARHGEF12* demonstrated RT Stop Scores less than -2, while weaker rG4s *EWSR1* and *HNRNPA2B1* had scores between -2 and 0. *PABPN1*, the non-rG4 control, had positives scores suggesting unfolded RNA (**Fig. 3C**).

**Figure 3.**
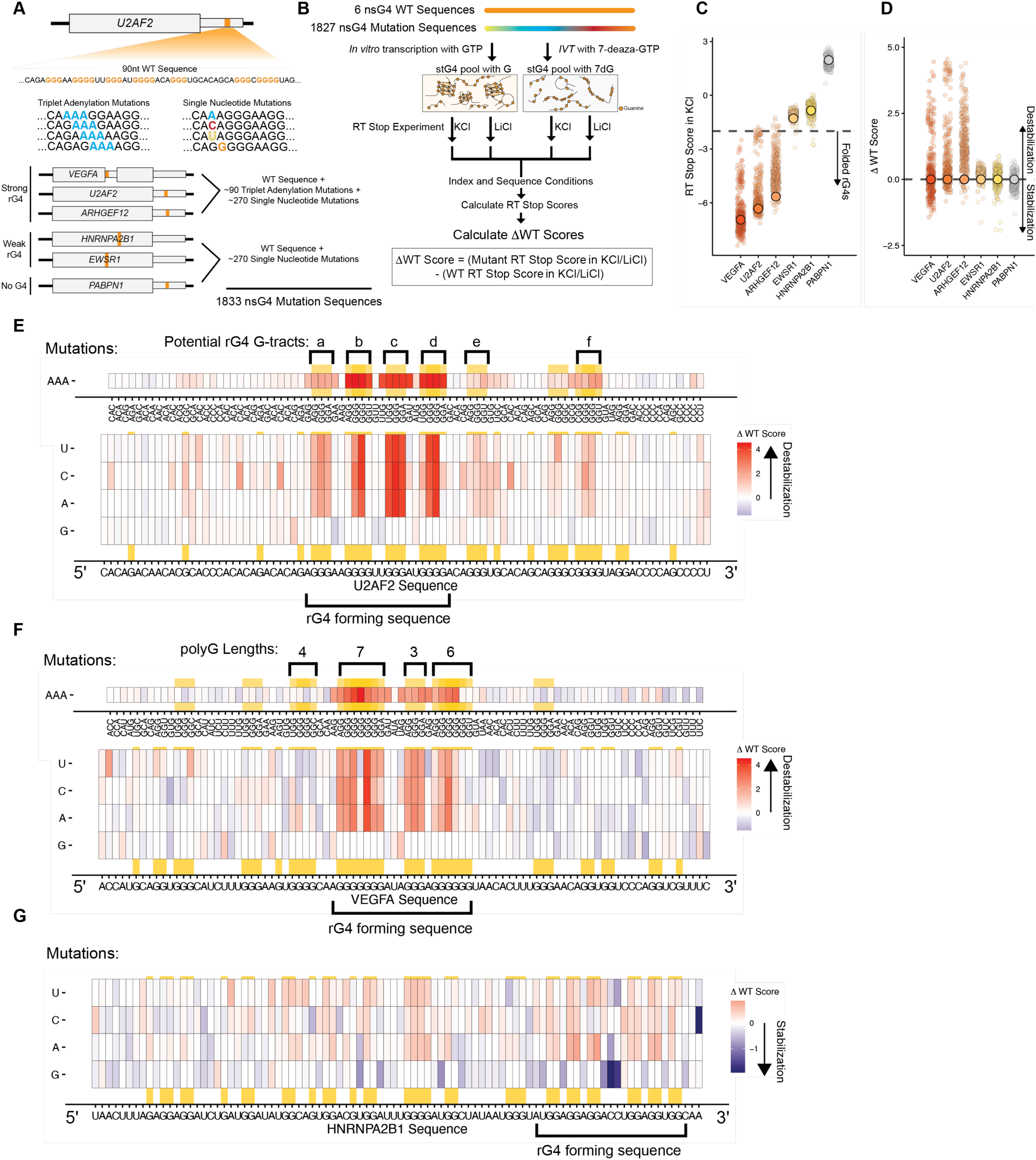
Mutational analysis of natural sequence rG4s. **A)** Schematic for the design of the mutational natural rG4 portion of the stG4 pool. *U2AF2* is used as an example, as the 90 nt sequence containing a rG4-forming region was tiled with either triplet adenylation (NNN > AAA) or single nucleotide (N > A, C, G, U) mutations. Three ‘strong’ natural rG4s were selected from *VEGFA*, *U2AF2*, and *ARHGEF12*, two ‘weak’ rG4s were selected from *HNRNPA2B1* and *EWSR1*, and one with no rG4 predicted from *PABPN1*. Orange rectangles show approximate location of the natural rG4 (3′UTR, coding sequence, or splice site), and the number of sequences from each natural rG4 with the mutational approaches shown. **B)** Schematic of RT stop sequencing procedure and evaluation of ΔWT Scores. Scatterplots of the natural sequence rG4s by **C)** RT Stop Scores in KCl and **D)** ΔWT Score (in KCl). WT sequence RT Stop Scores are shown as large circles colored by natural rG4, specifically *VEGFA* (red), *U2AF2* (dark orange), *ARHGEF12* (orange), *EWSR1* (light orange), *HNRNPA2B1* (yellow), and *PABPN1* (grey) (Data is the average of 3 independent replicates). Mutated sequences for each rG4 are shown as smaller circles behind the WT sequence with the same colors. Dashed horizontal lines show in **C)** -2 RT Stop Score for folded rG4s and **D)** 0 ΔWT Score. Mutation heatmaps of **E)** *U2AF2*, **F)** *VEGFA*, **G)** *HNRNPA2B1* rG4 sequences by ΔWT Score on a gradient from red (positive, destabilizing), white (0, equal to WT), to blue (negative, stabilizing) (Data is the average of 3 independent replicates). Sequences were plotted by position mutated, and the full WT sequence for the natural rG4 is shown (x-axis, bottom heatmap). Top heatmap (for **E** and **G** only) shows impact of triplet mutations to AAA, while the bottom heatmap shows single nucleotide mutations (U, C, A, G) at each position. Triplets which contain GGG and their directly adjacent base triplets (*top*) are colored in orange, while single G nucleotides (*bottom*) are colored in orange. *U2AF2* heatmap is annotated with potential G-tracts (brackets on top, lettered *a, b, c, d, e, f*), *VEGFA* heatmap is annotated with polyG sequence lengths (brackets on top, numbered). Natural sequence rG4s are annotated by predicted rG4 forming sequence (bracket on bottom).

To assess sequence features governing folding of these natural rG4s, we tiled all possible single nucleotide mutations as well as triplet A substitutions in single nucleotide steps along the full sequences (**Fig. 3A**). These mutants offered the ability to understand how mutations inside and outside of the predicted rG4 impact folding. To assay these natural RNA sequences, we derived a ΔWT score. Using the previously defined RT Stop Score metric, we calculated the difference between the RT Stop Score of the mutated sequence and the WT sequence (**Fig. 3B**). This served as a simple measure of the impact of a given mutation on RT stops. Positive ΔWT Scores indicated rG4 destabilization while negative scores indicated stabilization.

*U2AF2 reveals complex rG4 conformational selection:* We assessed ΔWT Scores along the sequence for AAA mutations and single nucleotide mutations (**Fig. 3E**). As noted in the highlighted regions, the *U2AF2* sequence contains 6 potential rG4 forming G-tracts. Our mutational scan identified three central G-tracts (**Fig. 3E**, *G-tracts b, c, d*) that, if individually mutated, led to significant destabilization. Three other G-tracts exhibit more limited destabilization phenotypes when mutated (**Fig. 3E**, *G-tracts a, e, f*), which may be due to their transient participation in G4 folding. The relative importance of each G-tract was further evaluated by single point mutations (**Fig. 3E**, *bottom*), in which we found an agreement with triplet mutations both in position and, surprisingly, magnitude of destabilization. Further analysis of flanking sequences revealed that mutation to Cs, especially in positions that generate CCC (*e.g.*, CAC->CCC) also destabilized the rG4 fold, presumably due to increasing the likelihood of CCC pairing with a GGG G-tract. This finding highlights the complexity of canonical base pairing influencing rG4 formation.

To better understand these effects, we modeled the *U2AF2* rG4 structure using *in silico* RNA folding. We found that G-tract *e* was involved in a long-range stem helix with the rG4 forming sequences in between, implying that this conformation contributes to rG4 folding or stabilization (**Fig. S3A**). G-tract *f* was found to be base-pairing with downstream C-rich sequences, and loss of *f* appeared to free a CCC patch, thus destabilizing the rG4 (**Fig. S3A**). Another potential mechanism is that hairpin formation stabilizes G4 folding—a feature well described in DNA G4s but underexplored for rG4s. More broadly, structural modeling revealed that ΔWT scores correlate with predicted minimum free energy (ΔΔG) of folding between rG4-containing and non-rG4-containing structures (**Fig. S3A**). Therefore, these analyses further highlight the complex interplay between base-pairing and rG4 formation, providing *in silico* evidence to support experimental evidence.

*VEGFA demonstrates favorability of discrete conformations:* The sequence of this rG4 contains multiple long polyG sequences, potentially supporting multiple rG4 folding topologies. However, single point mutations within the *VEGFA* sequence revealed critical rG4 forming GGG G-tracts within the sequence (**Fig. 3F**). Mutagenesis clearly defined 4 tracts within the G-rich stretch (<u>GGG</u>G<u>GGG</u>AUA<u>GGG</u>A<u>GGG</u>GGG) and showed that the last 3 guanines of this stretch were dispensable (**Fig. 3F**). Interestingly, mutations of Gs to Us induced stabilization both inside and outside of the defined rG4 forming region (**Fig. 3F**). For example, introduction of a U in the middle of the annotated 7 nt polyG sequence (GGG<u>G</u>GGG to GGG<u>U</u>GGG), had a stabilizing effect, mirroring our findings from above where U loops improved rG4 stability. Several critical findings emerged for this particular rG4, including: **i**) long G-runs were disfavored over short distinct G-tracts with one nucleotide loops (particularly a U loop), and **ii**) polyG-runs outside of the core rG4 forming sequence diminished rG4 folding, as mutating them enhanced stability. We propose that well-defined and spaced G-tracts improve stability, perhaps due to limiting dynamics between alternatively folded states.

*HNRNPA2B1 shows propensity for mutationally-improved rG4 folding:* The weak *HNRNPA2B1* rG4 sequence features numerous GG G-tracts, which complicate the identification of specific G-tracts involved in rG4 folding. Mutational analysis revealed few positions with a clear destabilization effect, suggesting that this sequence lacks a single, dominant rG4 structure. However, there were single nucleotide mutations which demonstrated strong stabilization effects (< -1 ΔWT Score) (**Fig. 3G**). Our attention was drawn to the annotated rG4 forming sequence which contains six GG G-tracts separated by primarily one nucleotide loops and a central CC (<u>GG</u>A<u>GG</u>A<u>GG</u>ACCU<u>GG</u>A<u>GG</u>U<u>GG</u>) (**Fig. 3G**). While destabilization was observed in mutations of the GG (noted in the rG4 forming sequence), a more striking stabilizing effect was observed upon mutation of the central CC residues, especially those with mutations from C to G (**Fig. 3G**). This suggests competition of canonical base-pairing with rG4 formation, as removal of these Cs ultimately led to improved rG4 stability.

Mutations in *ARHGEF12* (**Fig. S3B**), *EWSR1* (**Fig. S3C**), and *PABPN1* (**Fig. S3D**) sequences recapitulated many of the trends discussed above. Importantly, mutagenic analysis in the presence of LiCl rather than KCl also revealed minimal to no impact of mutations on RT stop score (ΔWT ∼ 0; **Fig. S3E, F**), confirming that the ΔWT scores directly reported on changes in rG4 folding stability. Overall, these mutagenesis experiments highlight the complexity and contributions of individual nucleotides to natural rG4 sequences.

### PDS shows stabilization of both longer loop GGGG-rG4s and mixed rG4s with U-rich loops

Many small molecule rG4 ligands have been identified and shown to alter stability and function of rG4s (41,42,53,62). However, systematically assessing the specificity of ligand-mediated stabilization towards different rG4s is a challenging problem that requires testing many sequences. We used our stG4 pool to evaluate the preferences of pyridostatin (PDS), one of the first small molecules to target G4s and used for global G4 stabilization (**Fig. 4A, B**) (54,55,63).

**Figure 4.**
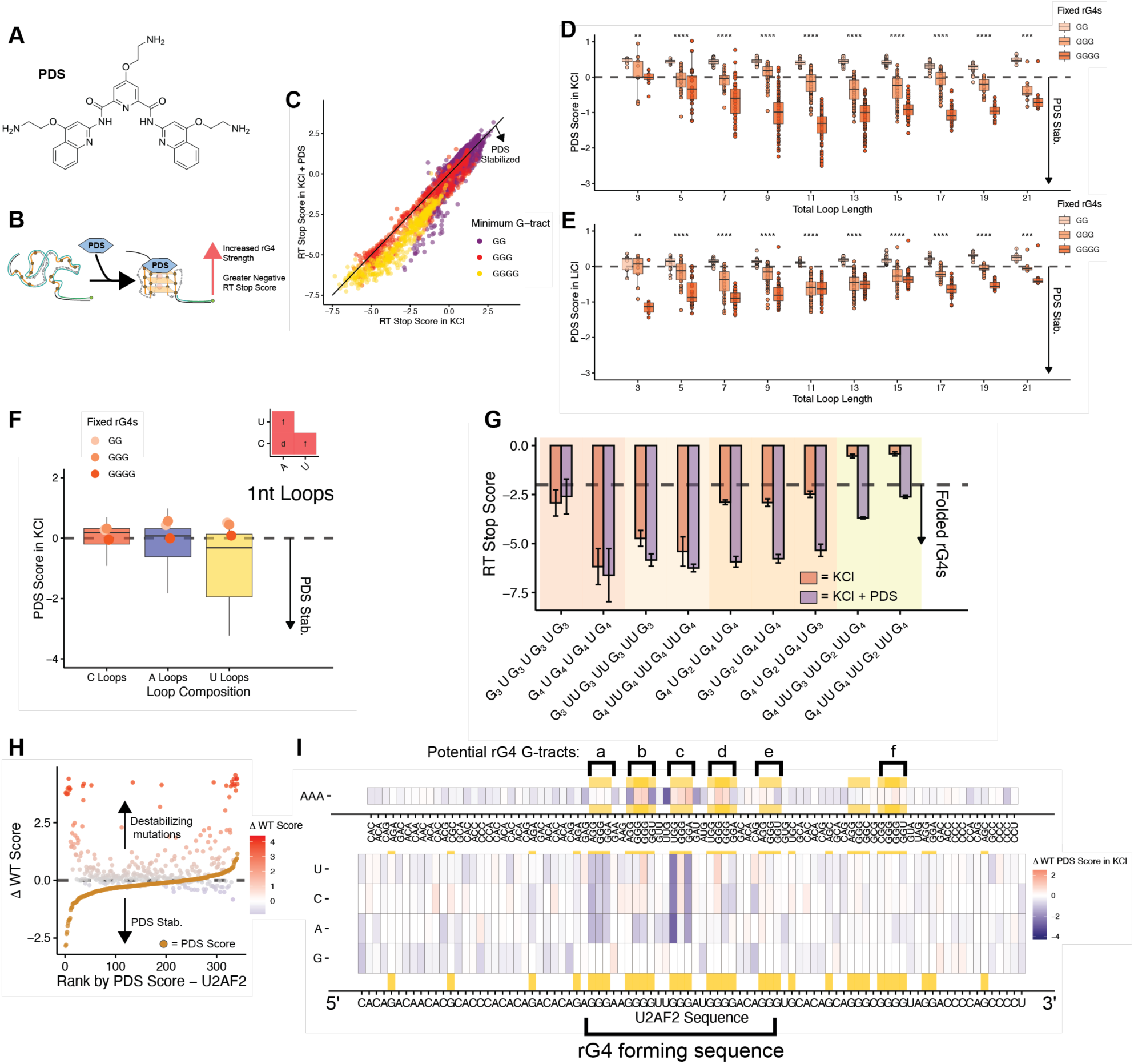
PDS stabilization is driven by unique structural characteristics. **A)** Chemical structure of pyridostatin (PDS). **B)** Schematic of expected interaction and RT Stop consequence by PDS addition. **C)** Scatter plots of putative rG4 sequences by RT Stop Score in KCl and PDS (y-axis) v. RT Stop Score in KCl alone (x-axis) colored by minimum G-tract length (GG (purple), GGG (red), GGGG (yellow)) (Data is the average of 3 independent replicates). Dashed horizontal line shows PDS score of 0. **D)** Box plot of putative rG4s by PDS Score in KCl with total loop length. Colored by fixed G-tract length (GG (light orange), GGG (orange), GGGG (dark orange)) (Data is the average of 3 independent replicates). Dashed horizontal line shows PDS score of 0. **E)** Box plot of putative rG4s by PDS Score in LiCl (log_2_(Sequence Freq_LiCl + PDS_/Sequence Freq_LiCl_) with total loop length. Colored by fixed G-tract length (GG (light orange), GGG (orange), GGGG (dark orange)) (Data is the average of 3 independent replicates). Dashed horizontal line shows PDS score of 0. **F)** Box plot of putative rG4s by PDS Score with different loop compositions (C (red), A (blue), U (gold)). Fixed G-tracts rG4s are shown as points colored by their G-tract length (GG (light orange), GGG (orange), GGGG (dark orange)). Dashed horizontal line shows PDS score of 0. Inset shows p-values determined by two-sided KS test corrected by BH procedure. Red square indicates p≤0.05. Values are as follows: d (p≤0.01), f (p≤0.0001). **G)** Bar graphs of putative rG4s with U loops and their respective average RT Stop Scores in KCl (red) and in KCl and PDS (purple) (n=3). Error bars (SD) generated from three independent replicates. Dashed line at -2 RT Stop Score is indicative for folded rG4s. Colored rectangles cluster sequences by observed trends by PDS-induced decreases in RT Stop Scores, either: no change (red), slight decrease (light orange), decrease (orange), and large decrease changing to ‘folded rG4’ (yellow). **H)** Ranked plot of *U2AF2* mutation sequences which correlate sequence ΔWT Score (colors correspond to ΔWT Score (red is positive, destabilizing; grey is zero; blue is negative, stabilizing)) and PDS Score (orange) (PDS score plotted on same axis) (Data is the average of 3 independent replicates). Dashed horizontal line shows ΔWT Score and PDS Score of 0. **I)** Combined heatmap of the *U2AF2* rG4 sequences by ΔWT PDS Score. Fill gradient shifts from red (positive, more readthrough in PDS), white (0, equal to WT), to blue (negative, less readthrough in PDS) were used (Data is the average of 3 independent replicates). Sequences were plotted by nucleotide or triplet mutated and full WT sequence for the natural rG4 is shown (x-axis, *bottom* heatmap). Triplets which contain GGG and their directly adjacent triplets (*top*) are colored in orange, while single G nucleotides (*bottom*) are colored in orange.

RT stop profiling revealed that addition of PDS triggered stabilization of putative rG4s while little effect was observed on non-rG4 controls (**Fig. S4A**). To investigate PDS-mediated stabilization, sequences were given a ‘PDS Score’, defined as the difference in RT Stop Scores with and without PDS, with more negative scores reflecting increased stabilization (**Fig. S4B**). Analysis of these sequences revealed three main phenomena of PDS stabilization: **i**) PDS generally stabilized rG4s both in KCl and LiCl as noted by more negative RT Stop Scores with PDS, specifically for the putative rG4 sequences (**Fig. 4C, Fig. S4C**). GGGG-containing rG4s, which were only weakly stable in LiCl, exhibited a high degree of stabilization with addition of PDS. **ii**) Fixed GGG- and GGGG- rG4s with longer destabilizing loops were overall stabilized, indicating that if an RNA has features that both enhance stability (*e.g.,* longer G-tracts) and decrease stability (longer loops), PDS can stabilize the rG4, regardless of the available cations (**Fig. 4D, E**). However, GG-rG4s, which were already weak, exhibited minimal PDS stabilization. **iii**) Mixed G-tract rG4s, especially those with a single GG G-tract, were uniquely stabilized (**Fig. 4C**). Fixed rG4s with short loops (3 nt total loops) did not change upon PDS addition in KCl (**Fig. 4D, F**). However, mixed rG4s with these short loops were stabilized upon PDS addition, in particular when loops contained Us (**Fig. 4F, Fig. S4D, E**). Closer examination of these mixed rG4s identified that rG4s which follow the motif G_4_U**G_<4_**U**G_<4_**UG_4_, with a single GG G-tract either in the G2 or G3 positions, were most strongly stabilized by PDS addition (**Fig. 4G**).

We next determined PDS-induced stabilization of our natural sequence rG4s, providing insight into how rG4-targeting drugs may be able to modulate folding of rG4s impacted by point mutations. The *VEGFA* rG4 demonstrated that PDS stabilization overcame the most destabilizing AAA and single point mutations (**Fig. S4F, G**). On the other hand, *U2AF2* showed a more complex pattern of stabilization, with some strongly destabilizing mutations being PDS-stabilized, but many other mutations not being stabilized through PDS addition (**Fig. 4H**). Mutations to the central G-tract of *U2AF2* (G-tract *c*, **Fig. 4H**) that generate a mixed G-tract rG4 were significantly stabilized by PDS (**Fig. 4I**). Thus, the mechanism of mixed G-tract rG4 stabilization by PDS described above is also observed for this natural sequence rG4. The weaker *HNRNPA2B1* rG4 displayed stabilization by addition of PDS, and the effect of some stabilizing mutations, specifically those that generated longer G-tracts (GG to GGG) were amplified by small molecule addition (**Fig. S4H, I**). Together, these analyses indicate that PDS recognition and stabilization is rather complex and involves both G-tract features, loop lengths, loop composition, and the initial stability of the rG4 in the absence of PDS.

### RNA Bind-n-Seq reveals rG4 preferences for FMRP LCD and G3BP1

rG4 function is intertwined with folding stability and ability to bind RBPs. A significant limitation in the field has been a lack of understanding of how rG4 features beyond the core fold, such as loop length and composition, impact RBP binding. To address this question, we purified two known rG4-binding RBPs, including full-length G3BP1 and the low complexity domain (LCD) of FMRP (**Fig. 5A**). To characterize interactions between these RBPs and rG4s, we employed RNA Bind-n-Seq (RBNS) (56,64), a large-scale binding assay that captures and quantifies RNAs that bind a given protein by deep sequencing (**Fig. 5B**, **Methods**). Recombinant proteins were incubated with the stG4 pool generated with either guanine or 7dG to assess the contributions by both rG4 structure and G-rich sequences towards the RBP-rG4 binding interaction (**Fig. 5B**). Upon isolating and sequencing the protein-bound RNA sequences, enrichment (*R*) values were derived by comparing the frequency of the protein-bound RNA sequences to the frequency of the RNA sequence in the guanine or 7dG pool inputs. A higher *R* value therefore indicates stronger binding. However, in this binding assay format RT of bound RNAs was performed in LiCl to limit RT stops to enable an accurate measure of binding strength.

**Figure 5.**
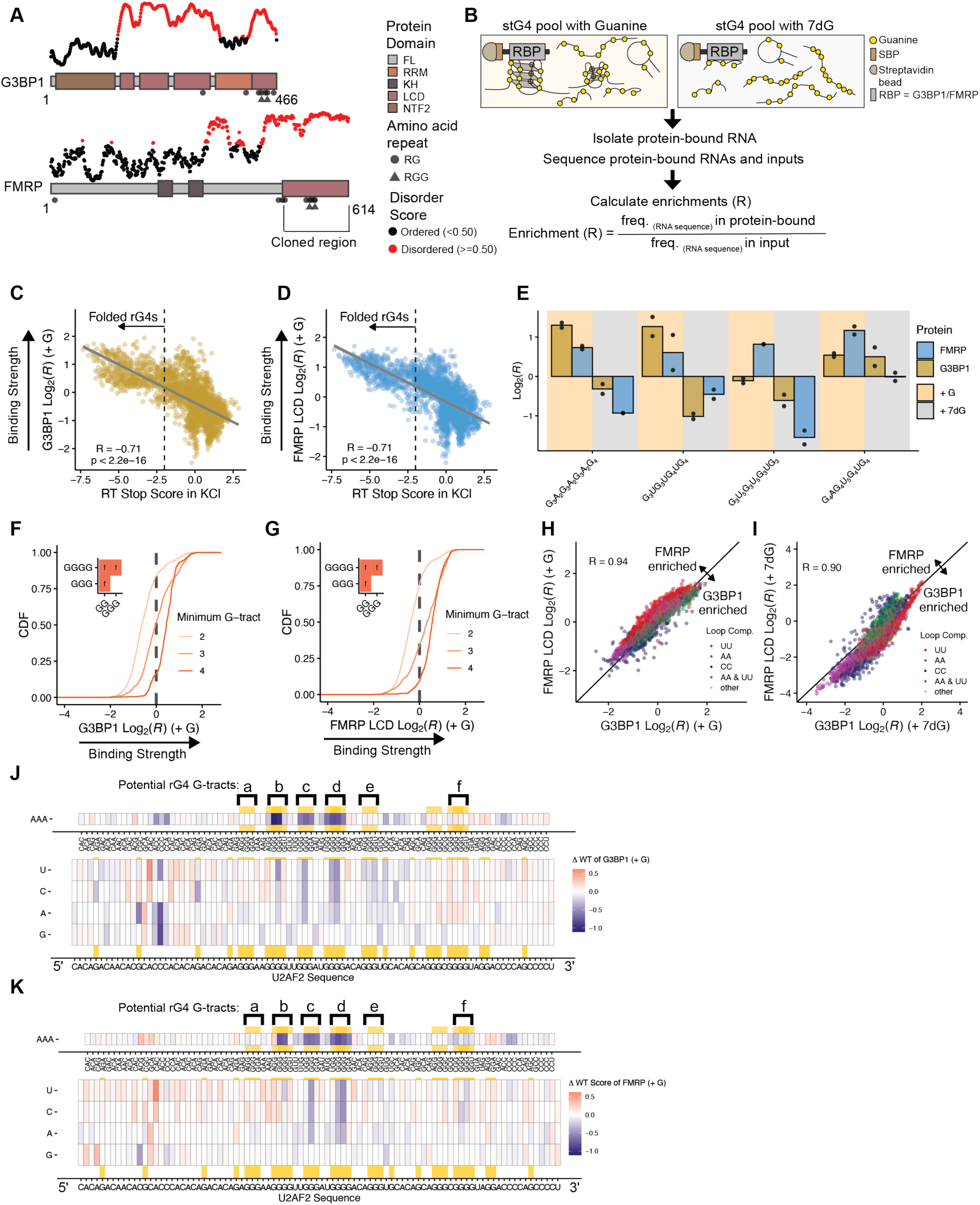
Analysis of protein binding preferences towards rG4s. **A)** Schematic of G3BP1 and FMRP with their respective disorder propensity plot. **B)** Schematic of RNA bind-n-seq (RBNS). Correlation plots of the log_2_ RBNS enrichment of **C)** G3BP1 (+ Guanine) (y- axis) and **D)** FMRP LCD (+ Guanine) (y-axis) versus RT Stop Score in KCl (x-axis). Pearson’s correlation coefficient and p-value are included. **E)** Log_2_ RBNS enrichment (Data is the average of 2 independent replicates) of unique oligos bound by FMRP LCD (light blue) and G3BP1 (dark gold). Dark gold rectangles represent oligos in stG4 (+ Guanine) pool. Grey rectangles represent oligos in stG4 (+ 7dG) pool. CDF of log_2_ RBNS enrichment of **F)** G3BP1 (+ Guanine) and **G)** FMRP LCD (+ Guanine) separated by the minimum G-tract allowed in an oligo. Inset shows p-values determined by two-sided KS test corrected by BH procedure. Red square indicates p≤0.05. Values are as follows: f (p≤0.0001). Scatter plots of the log_2_ RBNS enrichment of FMRP LCD (y-axis) versus G3BP1 (x-axis) in **H)** (+ Guanine) and **I)** (**+** 7dG). Colors represent combinations of the loop motifs of the putative rG4s. Heatmap of *U2AF2* by ΔWT Score of **J)** G3BP1 and **K)** FMRP LCD. Gradient represents the log_2_ RBNS enrichments for the change in WT sequence versus mutated sequence. Sequences were plotted by nucleotide or triplet mutated (for **A** only), and full WT sequence for the natural rG4 is shown (x-axis, *bottom* heatmap). Top heatmap (for **A** only) shows mutations of triplets to AAA and their impact, while the bottom heatmap shows single nucleotide mutations (U, C, A, G) at each position. Triplets which contain GGG and their directly adjacent triplets (*top*) are colored in orange, while single G nucleotides (*bottom*) are colored in orange.

As a first pass, we sought to understand the correlation between rG4 folding strength and protein binding. Comparison of RT Stop Score to protein binding strengths (*R* values) revealed significant correlations for both G3BP1 and FMRP LCD (**Fig. 5C, D**), and demonstrated that stronger rG4s tracks with higher binding enrichments (**Fig. S5E, F**). This relationship was limited with 7dG RNAs (**Fig. S5G, H**), supporting that the rG4 structure is critical for RBP interactions. The diversity of our pool also enabled us to investigate rG4 binding preferences by G3BP1 and FMRP. For example, we observed sequences that bound better to G3BP1 than FMRP in a structure- dependent manner, as well as the inverse (**Fig. 5E**). This differential recognition appeared to be independent of G- tract length (**Fig. 5F, G**), but rather driven by loop composition, with U loops being preferred by FMRP (**Fig. 5H**). In addition, G3BP1 and FMRP binding were mostly dictated by rG4 folding, as substitution by 7dG reduced enrichments overall (**Fig. S5I, J**). However, loop composition analysis showed that when rG4 folding was inhibited, RNAs were better enriched to G3BP1, perhaps due to the presence of other RNA binding domains in G3BP1, such as the RNA recognition motifs which are known to bind single-stranded RNA (**Fig. 5I**). While RBNS sequence enrichments were strongly correlated between FMRP and G3BP1 (**Fig. 5H**), we observed a limited overlap in the top binders for these proteins (**Fig. S5K, L**), further highlighting the complexity of rG4 recognition and RBP specificity.

We also evaluated the impact of G3BP1 and FMRP binding to our mutagenized natural sequences (discussed in **Fig. 3**). These proteins bound strongly to strong natural rG4s (*U2AF2*, *VEGFA*, *ARHGEF12*) and markedly less so to weaker rG4s and non-rG4 oligos (*EWSR1*, *HNRNPA2B1*, and *PABPN1*) (**Fig. S5M, N**). Generally, mutations to the natural sequences led to little impact on RBP enrichment (**Fig. S5M, N**). The largest variance in RBP binding was observed for *U2AF2* mutants where substitution of G-tracts for triple A mutants were the most detrimental to binding, but the effect was limited for single point mutants (**Fig. 5J, K**). However, comparison of FMRP and G3BP1 profiles for *U2AF2* revealed a set of single point mutations in a polyC sequence that decreased binding for only G3BP1 to similar levels as mutations to the core G-tracts (**Fig. 5J**). Thus, this pool can be used to directly compare stability to protein binding, and to distinguish sequence features preferred by different RBPs.

### pUG Quadruplexes lack RT stops, but demonstrate enrichment for RBPs

As mentioned above, poly(GU) sequences have recently been shown to form unique rG4s able to modulate biological outcomes (65). We included 29 of these putative ‘pUG quadruplexes’ with different numbers of (GU) repeats along with some additional sequence characteristics. RT stop assay revealed that all sequences, regardless of repeat number, demonstrated positive RT Stop Scores. PDS addition did not stabilized rG4 folding which implies that these pUG quadruplexes may not inhibit RTase readthrough (**Fig. S6A**). However, pUG sequences demonstrated significant binding enrichments with G3BP1 and FMRP LCD in RNAs generated with guanine but not with 7dG, and these were positively correlated with repeat number up to 14.5 p(GU) repeats (**Fig. S6B, C**). Taken together, the data suggest that while pUGs may not trigger an RT Stop, these sequences bind to RBPs through structure-dependent interactions.

## DISSCUSION

While study of rG4s has substantially increased over the past decades, challenges in understanding features that drive rG4 formation and stability remain. The canonical G4 motif, (G_≥2_N_1-7_)_4_, allows for massive variation (15) making it difficult to tease out rG4 characteristics that drive stability and/or recognition by proteins or targeting ligands. Therefore, we created a pool of thousands of synthetic putative rG4s where we varied G-tract lengths, loop lengths, and loop compositions (**Fig. 1A**) to define sequence-level drivers of rG4 formation by RT Stop quantification. Although not to the scale of this study, previous studies have used circular dichroism, thermal melting, and other biophysical approaches to identify contributions of cations and sequences for DNA or RNA G4s (16,17,59,60). These studies made significant strides in understanding G4 folding, correlating G4 stability with longer G-tracts, shorter loops, and short A loops, largely recapitulated within the current study. Our initial assessment of RT Stop conditions also supported the necessity of KCl for RT Stops (**Fig. 1F**), and found that the only significant temperature-induced changes in stability were observed for the strongest rG4s (**Fig. S1D**), in line with previous thermal melting studies (16). However, our approach builds on and overcomes limitations of this previous work. First, our use of RT stop experiments combined with sequencing allowed for thousands of rG4s to be quantitatively profiled in parallel. Second, sequencing afforded unique analysis and quantitative comparisons between each sequence, providing unprecedented depth into how specific rG4 variations or mutations perform within each assay. Third, the RNA preparation allowed for simple substitution of guanine and 7dG nucleotides into the same sequences, promoting an ‘unfolded rG4’ state that provided a baseline frequency for control normalization. Fourth, the use of 7dG also enabled us to determine if protein binding was rG4 structure dependent.

We also performed saturation mutagenesis on natural rG4 sequences to discern how mutations both in and outside of the rG4 forming sequence impact folding and even RBP recognition (**Fig. 3**, **Fig. 5J, K**). Mutagenesis of natural sequences revealed that within these 90 nt sequences, single point mutants were sufficient to alter rG4 stability (**Fig. 3**). This approach was useful in showing how single nucleotide polymorphisms (SNPs) in rG4 forming sequences modulate rG4 folding, as our selected natural rG4s harbor many annotated SNPs. Indeed, previous studies have either predicted or demonstrated SNP-mediated changes in rG4 formation for a limited set of rG4s (12,52,66). Turcotte and co-workers assessed SNP impacts on small molecule recognition of a G4 from ⍺- synuclein (52) and showed that a mutated sequence was stabilized by PDS. This is remarkably similar to what we observed for our weak rG4, *HNRNPA2B1*, as slightly stabilizing mutations without ligand (**Fig. 3G**) were magnified upon PDS addition (**Fig. S4L**). The *HNRNPA2B1* rG4 was tiled with ∼40 identified SNPs (reported in project gnomAD (67)), including SNPs which we show were rG4-stabilizing (**Fig. 3G**). While the biological impact of many of these SNPs are yet to be identified, our framework should reliably measure the impact of any mutation on any rG4 sequence.

Analysis of synthetic G4 recognition has been approached in similar manners by other groups. For example, Ray and co-workers developed a custom DNA G4 microarray to test molecular recognition (68,69). They tested binding of small molecules and proteins to ∼178,000 ssDNA 60mers with G4-folding propensity, allowing evaluation of specific G4 characteristics. They do mirror findings that we demonstrate herein, particularly that mutations within G-tracts were the most impactful to binding recognition, and that PDS had a T-rich loop preference (68). However, while their DNA-centric approach shows that interactions change in a KCl-dependent manner, it lacked the ability to directly assess specific G4 stability changes. Although our approach similarly does not directly evaluate PDS binding, we argue that our approach’s potential to identify drivers of ‘functional selectivity’ for G4- targeting ligands represents a critical step towards characterizing potential therapeutics that not only bind rG4s but further modulate specific rG4 structure-function relationships.

Many RBPs have been identified as rG4-binding proteins, including G3BP1 and FMRP (7,70–73), two factors studied here. By assessing binding of both full-length G3BP1 and the LCD of FMRP for thousands of rG4 forming sequences in competition, we found that binding was largely driven by folded rG4 stability (**Fig. 5C, D**), yet there were distinct sequence preferences (**Fig. 5E-I**). We also observed enhanced enrichment for natural rG4s by G3BP1 compared to the FMRP LCD (**Fig. S5M, N**), and that mutations that ablated rG4 folding showed the strongest decreases in binding for G3BP1 (**Fig. 5J, K**). Interestingly, however, G3BP1 binding was also decreased by specific mutations outside of the rG4 forming sequence (**Fig. 5J**), providing evidence that additional RNA motifs outside of the rG4 fold may be critical for specific rG4-RBP interactions. The sequence and rG4 topological preferences for RBPs are only starting to be investigated, for example with the RBPs HNRNPA1 (74) and NCL (75,76), representing a critical need to further *in vitro* understanding of RBP-rG4 recognition to build towards *in cellulo* interactions.

Despite *in cellulo* folding of these structures being a source of significant debate (29), there is an increasing body of evidence suggesting that rG4s play important roles in biology, including dynamic folding in response to stress and other stimuli (23). Furthermore, rG4s can be targeted with pharmacophores to modulate biological outcomes in cells (38–40), demonstrating the structures’ relevance as potential drug targets. By developing this massively parallel framework for investigating rG4 folding relationships and molecular recognition of rG4s, we propose an *in vitro* foundation for elucidating biochemical forces governing rG4 biology and providing opportunities to explain and derive *in cellulo* functional impacts.

## METHODS

### Design and *in vitro* transcription of stG4 pool

*Design of Structural Pool:* The structural rG4 pool was designed iteratively through the use of R (v. 4.1.0) and packages dplyr (v. 1.1.4) and tidyr (v. 1.3.1). Using the base G4 sequence composition of **G1-L1-G2-L2-G3- L3-G4** where Gs are G-tracts and Ls are loops, a few criteria were used to limit sequences selected and improve the diversity of sequences which could be evaluated. For G-tract variation, only three types of G-tracts were selected: GG, GGG, GGGG. We varied G-tract length at positions G1-4 (81 unique combinations) and made loop compositions the same nucleotide across all three loops (A, C, or U) with lengths locked from 1 nt to 7 nts from L1-L3. This contributed 1701 unique sequences. The second portion of sequences were designed to evaluate loop length and composition, allowing loop composition to vary between polyA and polyU sequences either 1, 3, 5, or 7 nts in length. All G-tracts were locked at GG, GGG, or GGGG to allow direct evaluation of these loop differences. This contributed another 1512 unique sequences for a total of 3213 sequences.

The oligo pool was set at a length of 90nts to complement with other natural sequences in the pool (see below). To accommodate varied sequence lengths (11 to 37 nts), a randomized surrounding sequence not expected to base pair with the rG4 forming region was selected. The 5’ flanking sequence (5’-ttcctcaagtcctgtgctacacatcgtatgtaccgaatc-3’) and the 3’ flanking sequence (5’- aaagtattatgttacgccgttcggatctgcacttccagca-3’) were appended to each synthetic rG4 and trimmed iteratively from either the 5’-end for the 5’ flanking sequence or 3’-end for the 3’ flanking sequence to accommodate the progressively longer putative rG4s. Oligos were then designed with the necessary 5’ and 3’ adapters to facilitate PCR and downstream methodology, leaving final 132nt sequences for synthesis.

*Design of Natural Sequences and Mutational G4 Pool:* The mutational G4 pool was designed from five natural sequences (*ARHGEF12*, *U2AF2*, *VEGFA*, *HNRNPA2B1*, *EWSR1*, and *PABPN1*) (**Table S1**) with different predicted G-quadruplex strengths by RNAFold and some identified in previous RT Stop analysis by Guo and Bartel (29). These 90 nt sequences were selected to reflect different rG4 strengths. These WT sequences were iteratively tiled by either AAA mutations and/or single point mutations across the full sequence, accounting for a total of 1890 sequences. Importantly, sequences were removed due to not being unique sequences (i.e., AAA mutations matching singling nucleotide polymorphisms) leaving 1833 unique sequences.

*Design of pUGs Oligos:* For the putative pUG quadruplexes sequences, different repeat numbers of pGU were selected and appended to the same flanking sequence used above to directly measure pUG stability rather than improved stability from the surrounding sequences. 6-18 repeats, with 9-15 repeats split in half repeats, were used for the base putative pUG quadruplexes accounting for 19 sequences. An additional 10 sequences were designed with 10 to 12 repeats (split in halves) bookended by AAA sequences, then also interspersed by a singly AAA in the center of the repeats.

*Design of Negative Controls:* Negative controls were designed iteratively using R (v. 4.1.0) and eCLIP data from ENCODE. These were separated into two categories, AAA controls and natural sequence RNA binding protein (RBP) binding sites. AAA controls were designed using the code below to generate putative ‘A-quadruplex’ sequences, locking AAA and varying the loop compositions and length. Specifically, each loop could either be polyC or polyU, and 3, 5, and/or 7 nts in length. Similarly to the putative rG4s described above, the same flanking sequences were used with the same iterative cropping to lead to 90nt base sequences before the addition of adapters to reach the final 132 nt in length. A total of 216 AAA controls were designed. Natural sequence RBP binding site sequences were selected from eCLIP peaks from EWSR1, FAM120A, FMR1, FUS, G3BP1, HNRNP(A1, C, K, L), and SRSF1. These 90 nt sequences were first evaluated by RNAFold(61) to confirm lack of rG4 folding prediction, and were binned by G-percentage. 354 sequences were selected with less than 25% G, and 355 sequences between 25% and 46% G, for a total of 709 putative unfolded natural control sequences from known G-quadruplex binding protein binding sites.

*Synthesis and Pool Preparation:* Structural pool oligos, negative controls, mutational G4 pool, and pUG oligos were synthesized in tandem by Twist Biosciences (**Table S1**). Oligo pool was amplified by polymerase chain reaction and a T7 promoter sequence was added with primers. *In vitro* transcription was performed using T7 Ribomax Express Large Scale RNA kit (Promega, #P1320) according to manufacturer protocols for the GTP structural pool. 7-deaza-GTP (Trilink Biotechnologies, N-1044) was substituted in the T7 Ribomax Large Scale RNA Production System (Promega, #P1280) at an equivalent concentration to the GTP for preparation of 7dG RNA pools and Strong rG4 sequences. After an overnight incubation, RNAs were subjected to a 1:1 phenol chloroform extraction followed by ethanol precipitation and polyacrylamide gel purification. RNAs were purified by a “crush and soak” protocol as gel slices were allowed to incubate overnight at 4°C with rotation in an RNA elution buffer (20 mM Tris-HCl, 250 mM NaOAc, 1 mM EDTA), before going through a second round of phenol chloroform and ethanol precipitation. Samples were resuspended in RNase-free water, then quantified by nanodrop and purity was confirmed by 6% TBE-Urea Novex (Invitrogen) gels. Two biological replicates were prepared from two separate PCR templates for the guanine and 7dG RNA pools.

### RT stop experiments

RT stop experiments were designed through inspiration from previous studies (77). RNA pools (guanine or 7dG) were first diluted to 3.75 nM with 3.75 nM of reverse transcriptase primer in RNase-free water. Solutions were then split into equivalent amounts dependent on the ion treatment (KCl, LiCl, or no salt), and 1/10 v/v was added of 1M (KCl, LiCl) or water for no salt. These were annealed at 100°C for 3 minutes, then slow-cooled on the benchtop for 10 minutes at room temperature. 10 µL of this solution was taken and added to a PCR strip for each RT reaction, and 2 µL of RNase-free water (or, for PDS (Sigma-Aldrich, SML2690) addition, 2 µL of 200 nM PDS in 0.2% DMSO) was added to each tube and allowed to incubate for 10 minutes at room temperature. 6 µL of this solution was added, and mixed by pipette, to 14 µL of the RT master mix which included 0.5 µL SuperScript III (Invitrogen, #18080), 1 µL 100 mM dNTPs (Promega, U1511), 1 µL 0.1 M DTT, 4 µL 5X In-house SuperScript III RT buffer with KCl/LiCl/No monovalent salt substitutions (250 mM Tris-HCl, 375 mM KCl/LiCl, 15 mM MgCl_2_, pH 8.3), and 7.5 µL of RNase-free water. Master mix was prepared per number of RT reactions per salt condition, and pre-incubated at 35°C for five minutes, then aliquoted into a PCR strip prior for the addition of 6 µL of annealed RNA pool and RT primer, leaving 20 µL total per RT reaction.

Temperature-dependent experiments were incubated at 30°C, 40°C, or 50°C for 40 minutes (run in duplicates). Otherwise, all other experiments were incubated at 40°C for 40 minutes (run in triplicates). All samples were heated for 5 minutes at 100°C, then RT samples were then prepared for sequencing by PCR amplification with Illumina indexes with between 10-13 PCR cycles (98°C for 2 minutes, (98°C for 30 seconds, 57°C for 20 seconds, 72°C for 30 seconds) for 10-12 cycles, then 72°C for 2 minutes. Products were evaluated by agarose gel electrophoresis, then purified through gel extraction and quantified by Qubit dsDNA kit (ThermoFisher, Q32851).

Libraries were sequenced using an Illumina NextSeq 1000. Initial RT condition tests were performed in duplicate, whereas other experiments were performed in triplicate. Mapping and counting the reads of each pool sequence per sample was performed using STAR. To avoid zero reads, each sequence was given one pseudocount prior to normalization. These read counts were then normalized by the total number of mapped reads in each sample, as a sequence frequency. RT Stop Score was calculated by dividing the frequency of a sequence with guanine by the frequency of the same sequence with 7dG in the same RT conditions. No read cutoff was instituted for the majority of the analyses in this work.

Gel experiments were performed similarly to as described above. The main modifications include changing the RNA concentrations for the specific ‘strong rG4’ 5’-(AGGG)_4_AAAAAAA-3’ with either guanine or 7dG to 40 nM and annealing with a FAM-labelled RT primer at 10 nM. Samples were annealed in either 1/10 v/v was added of 1M (KCl, LiCl), then the master mix was prepared similar to as described above with 0.5 µL of SuperScript III (Invitrogen, #18080), 1 µL dNTPs (Promega, U1511), 4 µL 5X In-house SuperScript III RT buffer with KCl/LiCl substituted (250 mM Tris-HCl, 375 mM KCl/LiCl, 15 mM MgCl_2_, pH 8.3), 1 µL of 0.1 M DTT, and 8.5 µL of RNase-free water. After the RT reaction, samples were treated with 3 µL of RNase A (ThermoScientific, EN0531) diluted 1:2 in water, and incubated for five minutes at 37°C. These samples were diluted 1:1 in 2X RNA dye (New England Biolabs, B0363S) and run on Novex™ 10% TBE-Urea Gels (Invitrogen, EC68795). These gels were then visualized on a Bio-Rad ChemiDoc Touch Imaging System using the Alexa488 channel and quantified using ImageJ.

### Protein purification of G3BP1 and FMRP

Full length (FL) human G3BP1 gene (Uniprot Q13283-1) and low complexity domain (LCD) of mouse FMRP (defined as residues 478–614, previously reported in (7)) were cloned into pGEX-6P-1. Plasmid were transformed into Rosetta *E. coli* cells (Millipore Sigma, 70954) and grown in LB media with 100 µg/mL ampicillin and 25 µg/mL chloramphenicol at 37°C until reaching an optical density of ∼0.8. Cell cultures were then cooled down to 4°C and induced with 0.5 mM IPTG overnight at 16°C. Cell cultures were pelleted by centrifugation at 4000 x g for 13 minutes at 4°C, supernatant was discarded, and pellets were resuspended with lysis buffer (1% Triton X-100, 5 mM DTT, 4 mM MgCl2, 200 mM NaCl, 20 mM HEPES, 1 tablet of protease inhibitor mini tablet, EDTA-free (Thermo Scientific #A32955) per 2 liters of culture). Lysate was sonicated and then incubated with 250 units of benzonase nuclease (Millipore #E1014) and 3 units of RQ1 RNAse-Free DNAse per liter of culture at 25°C for 15 minutes. Lysate was further clarified by centrifugation at 17,500 x g for 30 minutes and supernatant was collected.

G3BP1 FL and FMRP LCD were purified with glutathione agarose beads (Thermo Scientific, 16101). Beads equilibrated with lysis buffer were incubated with clarified cell lysate for 1 hour at 4°C followed by 3 washes with wash buffer (0.1% Triton X-100, 200 mM NaCl, 20 mM HEPES, 3.5 mM EDTA). G3BP1 was eluted for 1.5 hours at 25°C with cleavage buffer with PreScission Protease (10% glycerol, 5 mM DTT, 100 mM NaCl, 20 mM HEPES, 0.01% triton X-100, 1 mg/mL in-house PreScission Protease). Following, G3BP1 was further cleaned by heparin purification with a gradient from low salt buffer (50 mM NaCl, 50 mM HEPES, 3% glycerol) to high salt buffer (1 M NaCl, 50 mM HEPES, 3% glycerol) on a ÄTKA Pure HPLC, relevant fractions were pooled together, and Triton

X-100 was added for a final concentration of 0.01%. FMRP LCD was eluted for 1.5 hours at 25°C with elution buffer (20 mM GSH, 50 mM Tris Base pH 8.0) followed by dialysis in 150 mM NaCl and 20 mM HEPES. After dialysis, glycerol and Triton X-100 were added for a final concentration of 3% and 0.01%, respectively. Proteins were then concentrated with an Amicon Ultra 10 kDa centrifugal filter unit (#UFC8010) and concentration was determined by Pierce BCA Assay Kit (Thermo Scientific, #23208).

### RNA Bind-n-Seq (RBNS)

RBNS was performed as described in detail elsewhere(56,64). In brief, SBP-tagged G3BP1 FL and FMRP LCD were incubated with Dynabeads MyOne Streptavidin T1 (Invitrogen, #65602) for 30 minutes at 4°C in RBNS KCl binding buffer (25 mM Tris-HCl, 150 mM KCl, 3 mM MgCl_2_, 500 μg/mL BSA, 20 units/mL SUPERase IN (Invitrogen, AM2696)) or in RBNS LiCl binding buffer (25 mM Tris-HCl, 150 mM LiCl, 3 mM MgCl_2_, 500 μg/mL BSA, 20 units/mL SUPERase IN (Invitrogen, AM2696)). Prior to addition of the pools to binding reactions, stG4 and stG4 7dG pool were heated in either 150 mM KCl or 150 mM LiCl, respectively, at 100°C for 3 minutes followed by cool down at 25°C for at least 10 minutes. Protein-beads complexes were then incubated with either stG4 pool in RBNS KCl binding buffer or with stG4 7dG pool in RBNS LiCl binding buffer at a final concentration of 500 nM for 1 hour at 4°C. Beads with protein-RNA complexes were isolated in the magnetic stand (Invitrogen, 12321D) and thoroughly washed with either RBNS KCl wash buffer (25 mM Tris-HCl, 150 mM KCl, 20 units/mL SUPERase IN (Invitrogen, AM2696)) or with RBNS LiCl wash buffer (25 mM Tris-HCl, 150 mM LiCl, 20 units/mL SUPERase IN (Invitrogen, AM2696)). Protein-RNA complexes were then eluted twice with elution buffer (4 mM biotin, 25 mM Tris-HCl) for 30 minutes at 37°C. Eluates were pooled together and purified by phenol chloroform extraction. Prior to reverse transcription, purified protein-bound RNAs were heated to 100°C in 150 mM LiCl for 3 minutes and cooled down at 25°C for at least 10 minutes. Reverse transcription (RT) was performed with SuperScript III Reverse Transcriptase (Invitrogen, 18080044) and an RT primer (**Table S1**). A modification to the 5X-first strand buffer was made, where KCl was changed to LiCl, to prevent RNA structure. Libraries were amplified with RBNS index primers and sequenced using an Illumina NextSeq 1000.

## DATA ACCESSIBILITY

Raw FASTQ files, custom genomes and code used for analysis of the stG4 pool are available upon request.

## SUPPLEMENTAL DATA

Supplementary data is available online, including Supplemental Figures 1-6, Supplemental Table 1 (Sequences), and Supplemental Table 2 (Processed data for analysis).

## ACKNOWLEDGEMENTS

This work was in part supported by NIH T32 HL007149 (J.G.M.), 1T32GM135095-01 (B.B.G.), R35GM142864 (D.D.), R35GM147010 (A.M.), UNC Chapel Hill Simmons Scholar Award (M.M.A.), as well as startup funds from UNC Chapel Hill to (D.D.). The authors would also like to acknowledge G. A. Goda, A. S. Kavalipati, and J. M. Berger for critical reading and review of the manuscript.

## CONTRIBUTIONS

**Justin G. Martyr:** Writing – Original draft, Writing – Review and editing, Conceptualization, Formal analysis, Investigation, Methodology, Visualization. **Bryan B. Guzmán:** Writing – Original draft, Writing – Review and editing, Conceptualization, Formal analysis, Investigation, Methodology, Visualization. **Yue Hu:** Writing – Review and editing, Conceptualization, Visualization. **Rebekah L. Rothacher:** Writing – Review and editing, Visualization, Data curation. **Anthony M. Mustoe:** Writing – Review and editing, Conceptualization, Funding acquisition. **Maria M. Aleman:** Writing – Original draft, Writing – Review and editing, Funding acquisition. **Daniel Dominguez:** Writing – Original draft, Writing – Review and editing, Conceptualization, Investigation, Methodology, Funding acquisition, Supervision, Project administration, Data curation, Resources.

