## Supplemental Figures for "Massively parallel characterization of RNA G-quadruplex stability and molecular recognition"

[1]Department of Pharmacology, University of North Carolina, Chapel Hill, NC
[2]Blood Research Center, University of North Carolina, Chapel Hill, NC
[3]Therapeutic Innovation Center (THINC), Verna and Marrs McLean Department of Biochemistry and Molecular Pharmacology, Department of Molecular and Human Genetics, Baylor College of Medicine, Houston, TX
[4]McAllister Heart Institute, University of North Carolina, Chapel Hill, NC
[5]RNA Discovery Center, University of North Carolina, Chapel Hill, NC
[6]Department of Biochemistry and Biophysics, University of North Carolina, Chapel Hill, NC
[7]Bioinformatics and Computational Biology Curriculum, University of North Carolina, Chapel Hill, NC
[8]Lineberger Comprehensive Cancer Center, University of North Carolina, Chapel Hill, NC
*authors contributed equally

**Table of Contents:**


| | |
|---|---|
| **Figure S1. Temperature significantly impacts folded rG4s.** | **S-2** |
| **Figure S2. Additional rG4 characterization.** | **S-3** |
| **Figure S3. Additional mutational rG4s.** | **S-4,5** |
| **Figure S4. Additional PDS characterization.** | **S-6** |
| **Figure S5. Additional G3BP1 and FMRP LCD RBNS characterization.** | **S-7,8** |
| **Figure S6. pUGs lack RT stops but bind proteins.** | **S-9** |
| **Table S1: Sequences for pool oligos** | **.xlsx** |
| **Table S2: Processed RT Stop Scores and ΔWT Scores** | **.xlsx** |

**Supplemental Figure 1**

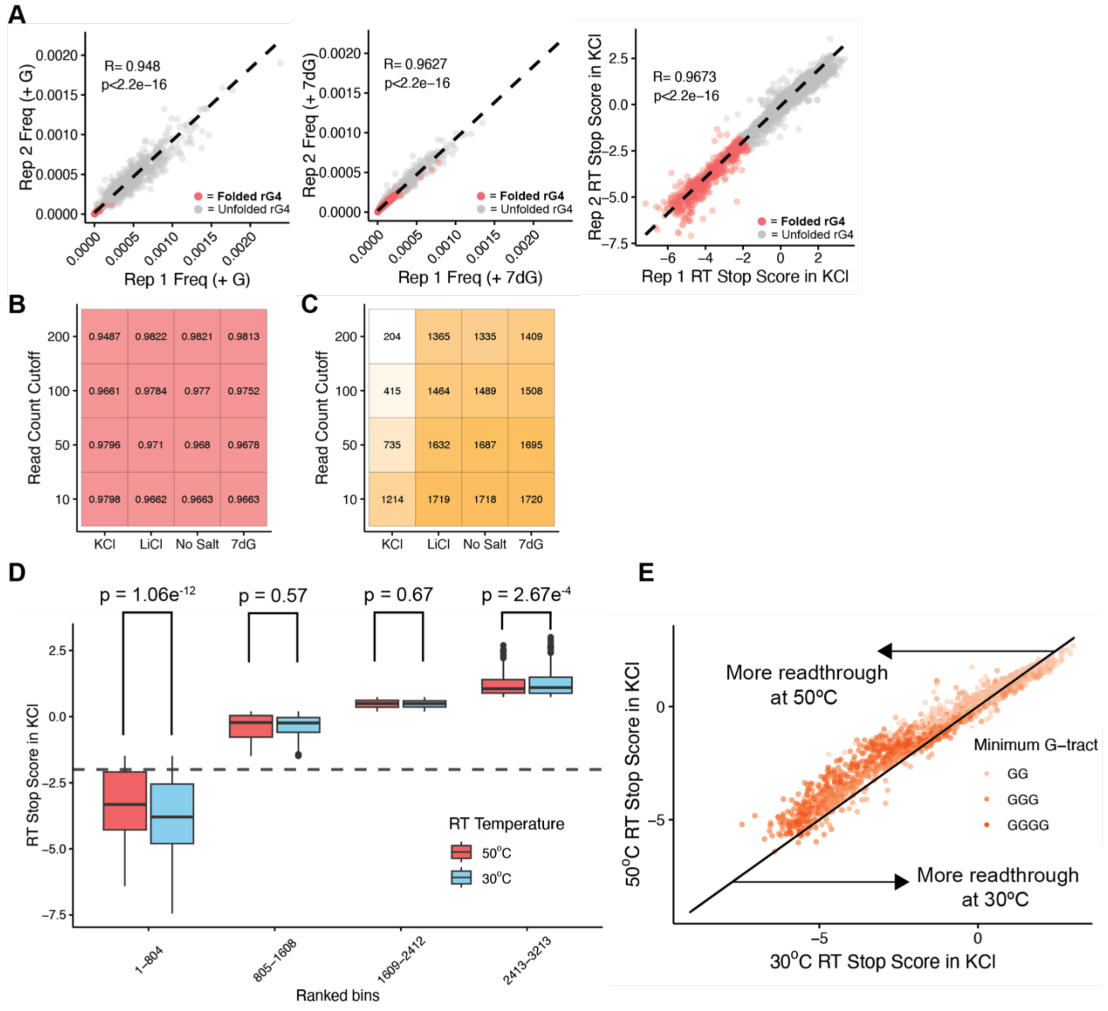


**Supplemental Figure 1: Temperature impacts on rG4 RT stops. A)** Correlation plots comparing (*left*) RNA frequencies between replicates of the rG4 structural pool with guanine in KCl, (*center*) RNA frequencies between replicates of the rG4 structural pool with 7dG in KCl, and (*right*) RT Stop Scores in KCl at 40ºC between replicates of the rG4 structural pool. Oligos that have an average RT Stop Score in KCl at -2 or below, identified as folded rG4s, are colored in red. Pearson's correlation coefficients (R values) and p-values are also shown. Matrix of replicate correlations of RT Stop Scores in KCl at 40ºC, with read cutoffs based on the number of reads per sequence in each specific treatment (KCl, LiCl, No Salt, or with 7dG in KCl) for **B)** Pearson's correlation coefficients (R values) and **C)** number of 'folded rG4s' counted. **D)** Box-plot analysis of putative rG4 sequences in equally-sized bins by ranked RT Stop Score in KCl comparing 30°C (blue) and 50°C (red) (Data is the average of 2 independent replicates). P-values were determined by student's two-sided t-test. **E)** Scatterplot of putative rG4 sequences based on their RT Stop Score in KCl at 30°C (x-axis) and 50°C (y-axis) (Data is the average of 2 independent replicates). Points are colored by minimum G-tract length (GG (light orange), GGG (orange), and GGGG (dark orange)) and black line represents a perfect correlation between RT Stop Scores. Above the line represents sequences that have improved readthrough at 50°C, and below the line are sequences with more readthrough at 30°C.

### Supplemental Figure 2

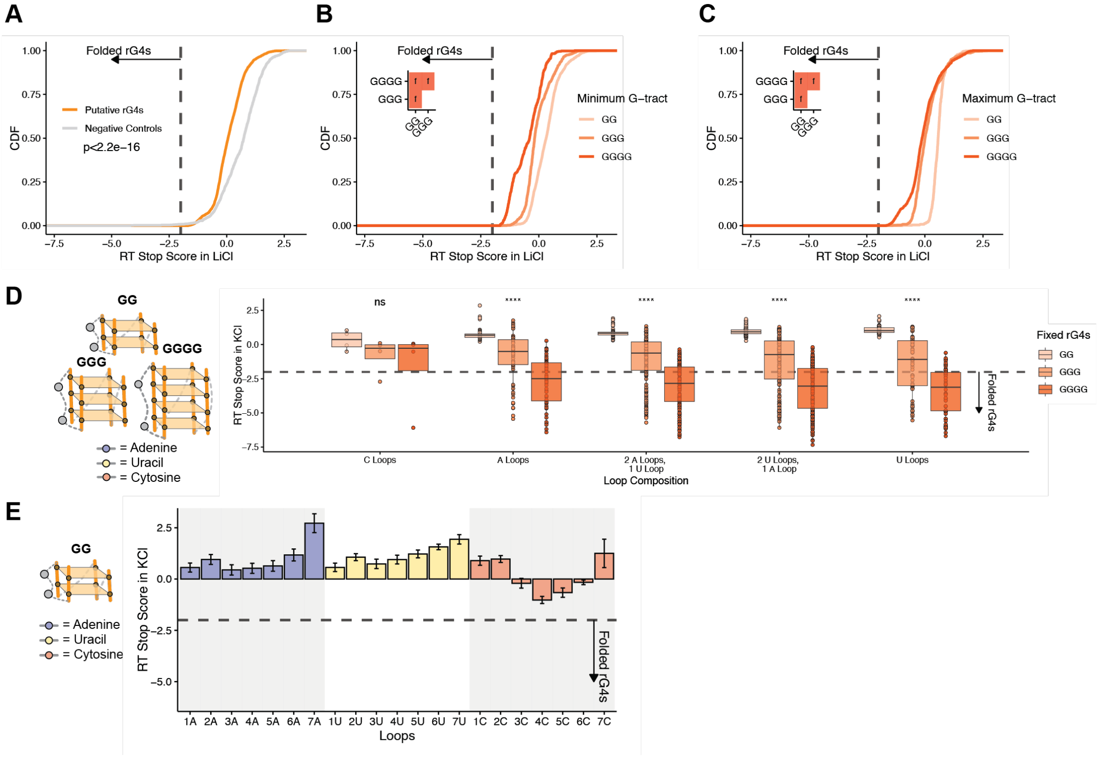


**Supplemental Figure 2: Additional rG4 characterization.** CDF of the structural rG4 pool oligo RT Stop Score in LiCl **A)** by putative rG4s (orange), and negative controls (grey) (Data is the average of 3 independent replicates), then further **B)** by minimum and **C)** by maximum G-tract length, specifically GG (light orange), GGG (orange), and GGGG (dark orange) (n=3). Line at -2 RT Stop Score is indicative of cutoff for folded rG4s. P-value is shown, or plot insets (*left*) show p-values determined by two-sided KS test corrected by BH procedure. Red square indicates $p \leq 0.05$. Values are as follows: f ($p \leq 0.0001$). **D)** G-quadruplex schematic (*left*) demonstrating selection of oligos possessing the fixed GG, GGG, and GGGG at all positions and varying the loop composition, either all C loops, all A loops, all U loops, or a combination of A and U loops. Box plot analysis of RT Stop Score in KCl (*right*) for oligos having fixed G-tract length of GG (light orange), GGG (orange), GGGG (dark orange) comparing loop composition (Data is the average of 3 independent replicates). Line at -2 RT Stop Score is indicative of cutoff for folded rG4s. P-values show significance of the distribution by minimum G-tract: ns = $p > 0.05$, **** = $p \leq 0.0001$. **E)** Bar plots for fixed GG-rG4s and the RT Stop Scores in KCl with differing loop lengths and compositions. This is 1 nt to 7 nt loops of the same nucleotide composition (i.e., 1A, 2A, 3A…) with A loops in blue, U loops in yellow, and C loops in red. Line at -2 RT Stop Score is indicative of cutoff for folded rG4s.

### Supplemental Figure 3

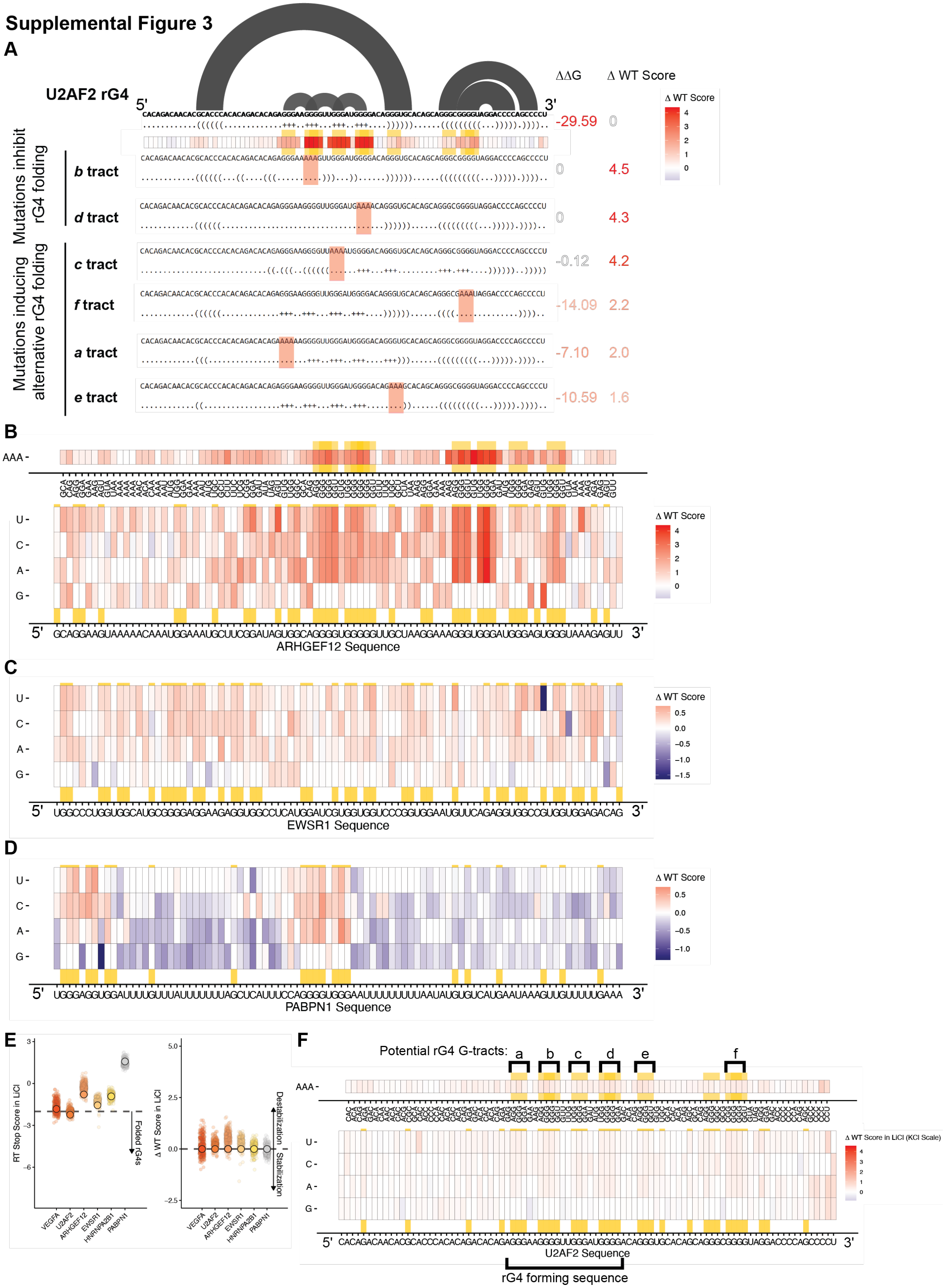

**Supplemental Figure 3: Additional mutational rG4s. A)** *In silico* RNA folding analysis of *U2AF2*. Mutations for specific tracts are shown with their impacts in folding shown by dot-bracket notation. ΔΔG was calculated by taking the predicted ΔG of folding when allowing G4 formation and subtracting the ΔG of folding when disallowing G4 formation (by RNAfold), with 0 meaning no G4 being predicted, and more negative ΔΔG meaning rG4 predicted. ΔWT scores are shown for *U2AF2*, demonstrating correlation between these metrics. Combined heatmaps of **B)** *ARHGEF12*, **C)** *EWSR1*, **D)** *PABPN1* sequences by ΔWT Score on a gradient from red (positive, destabilizing), white (0, equal to WT), to blue (negative, stabilizing) (Data is the average of 3 independent replicates). Sequences were plotted by nucleotide or triplet mutated (for **A** only), and full WT sequence for the natural rG4 is shown (x-axis, *bottom* heatmap). Top heatmap (for **A** only) shows mutations of triplets to AAA and their impact, while the bottom heatmap shows single nucleotide mutations (U, C, A, G) at each position. Triplets which contain GGG and their directly adjacent triplets (*top*) are colored in orange, while single G nucleotides (*bottom*) are colored in orange. **E)** Scatterplots of the natural sequence rG4s by RT Stop Score in LiCl and ΔWT Score (in LiCl). WT sequence RT-Stop Scores are shown as large circles colored by natural rG4, specifically *VEGFA* (red), *U2AF2* (dark orange), *ARHGEF12* (orange), *EWSR1* (light orange), *HNRNPA2B1* (yellow), and *PABPN1* (grey) (Data is the average of 3 independent replicates). Mutated sequences for each rG4 are shown as smaller circles behind the WT sequence with the same colors. Dashed horizontal lines show in (*right*) -2 RT Stop Score for folded rG4s and (*left*) 0 ΔWT Score. **F)** Combined heatmap of the *U2AF2* sequence by ΔWT Score in LiCl using theΔWT Score in KCl gradient from red (positive, destabilizing), white (0, equal to WT), to blue (negative, stabilizing) (Data is the average of 3 independent replicates). Sequences were plotted by nucleotide or triplet mutated and full WT sequence for the natural rG4 is shown (x-axis, bottom heatmap). Triplets which contain GGG and their directly adjacent triplets (*top*) are colored in yellow, while single G nucleotides (*bottom*) are colored in yellow.

### Supplemental Figure 4

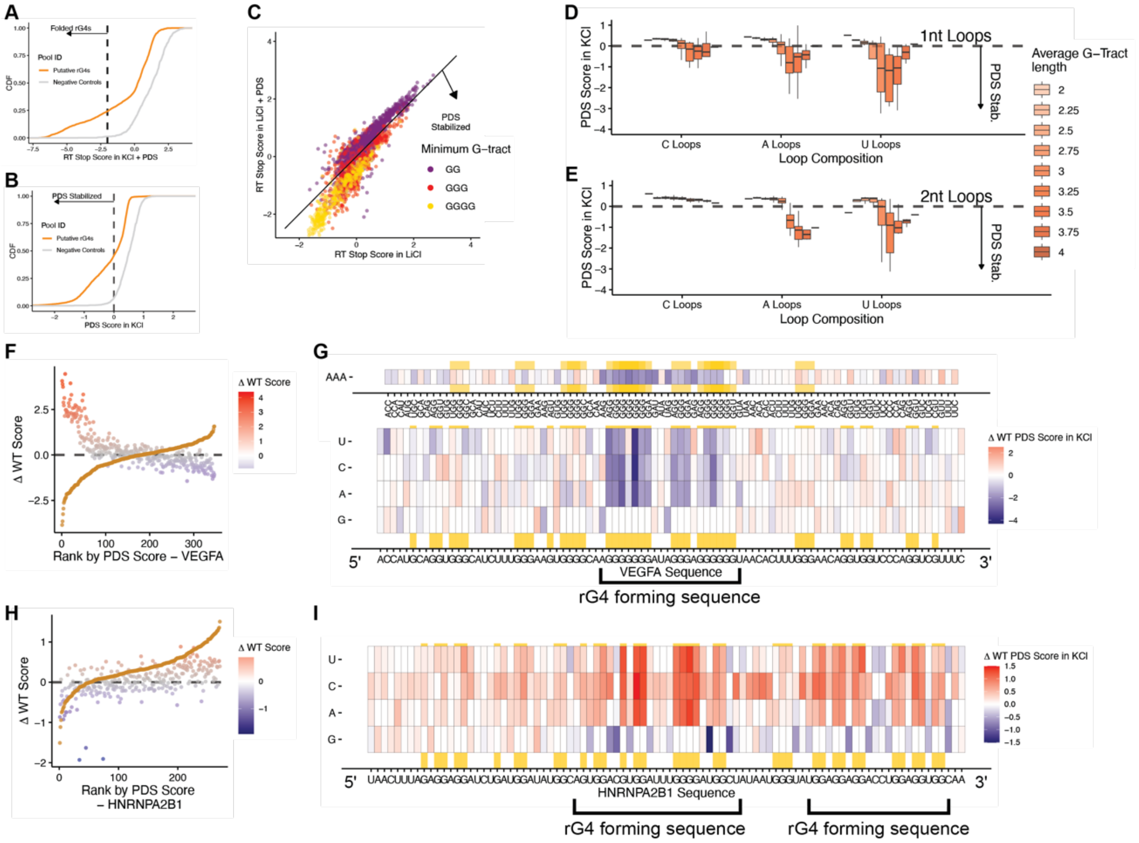


**Supplemental Figure 4. Additional PDS characterization. A)** CDF of RT Stop Scores in KCl with PDS by pool identity (putative rG4s (orange) and negative controls (grey)). Dashed vertical line at -2 RT Stop Score is indicative of cutoff for folded rG4s. **B)** CDF of PDS Scores in KCl by pool identity (putative rG4s (orange) and negative controls (grey)). Dashed vertical line shows PDS Score of 0. **C)** Scatter plots of putative rG4 sequences by RT Stop Score in LiCl and PDS (y-axis) v. RT Stop Score in LiCl (x-axis) colored by minimum G-tract length (GG (purple), GGG (red), GGGG (yellow)) (Data is the average of 3 independent replicates). Sequences to the right of the line are PDS stabilized. Box plots demonstrating distribution of PDS Score in KCl for different loop compositions (C, A, U) with different average G-tract lengths (gradient of light orange for all GG (2), dark orange for all GGGG (4)) for **D)** 1 nt and **E)** 2 nt loops. PDS score less than 0 corresponds to increased rG4 stability and decreased readthrough. Ranked plots of mutation library sequences by natural rG4 which correlate sequence ΔWT Score (colors correspond to ΔWT Score (red is positive, destabilizing; grey is zero; blue is negative, stabilizing)) and PDS Score (orange) for **F)** *VEGFA*, and **H)** *HNRNPA2B1*. Dashed lines at a ΔWT and PDS Score of 0 corresponds to those sequences that are stabilized below the line or destabilized above the line. Combined heatmaps of the **G)** *VEGFA* and **I)** *HNRNPA2B1* rG4 sequences by ΔWT PDS Score. A gradient from red (positive, destabilizing in PDS), white (0, equal to WT), to blue (negative, stabilizing in PDS) was used and independently set for each rG4 (Data is the average of 3 independent replicates). Sequences were plotted by nucleotide or triplet mutated and full WT sequence for the natural rG4 is shown (x-axis, *bottom* heatmap). Triplets which contain GGG and their directly adjacent triplets (*top*) are colored in orange, while single G nucleotides (*bottom*) are colored in orange.

### Supplemental Figure 5

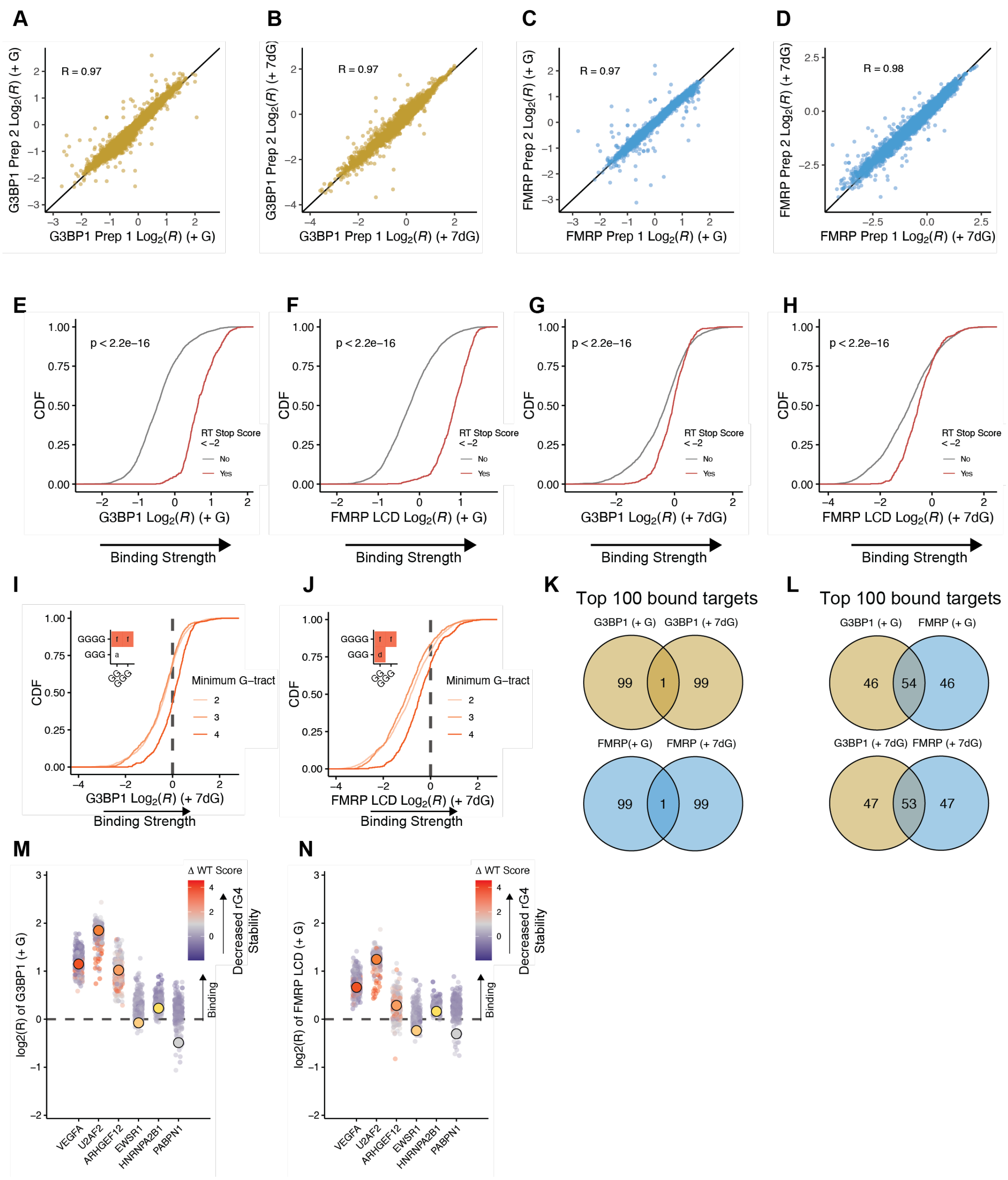


**Supplemental Figure 5. Additional G3BP1 and FMRP LCD RBNS characterization.** Correlation plots comparing the $\log_2$ RBNS enrichment across protein preps (Data is the average of 2 independent replicates) for **A)** G3BP1 (+ Guanine), **B)** G3BP1 (+ 7dG), **C)** FMRP LCD (+ Guanine), **D)** FMRP LCD (+ 7dG). CDF of $\log_2$ RBNS enrichment of **E)** G3BP1 (+ Guanine), **F)** FMRP LCD (+ Guanine), **G)** G3BP1 (+ 7dG) and **H)** FMRP LCD (+ 7dG) separated by RT Stop Score (Yes or No). RT Stop

Score – Yes are oligos that had a RT Stop Score in KCl at -2 or below. RT Stop Score – No are oligos that had a RT Stop Score in KCl greater than -2. P-value was calculated via two-sided KS test. CDF of $log_2$ RBNS enrichment of **I)** G3BP1 (+ 7dG) and **J)** FMRP LCD (+ 7dG) separated by the minimum G-tract allowed in an oligo. Inset show p-values determined by two-sided KS test corrected by BH procedure. Red square indicates $p \leq 0.05$. Values are as follows: a (ns), d ($p \leq 0.01$), f ($p \leq 0.0001$). **K)** Venn diagram of the top 100 bound targets based on $log_2$ RBNS enrichment for (*top*) G3BP1 (+ Guanine) and G3BP1 (+ 7dG) and (*bottom*) FMRP LCD (+ Guanine) and FMRP LCD (+ 7dG). **L)** Venn diagram of the top 100 bound targets based on $log_2$ RBNS enrichment for (*top*) G3BP1 (+ Guanine) and FMRP LCD (+ Guanine) and (*bottom*) G3BP1 (+ 7dG) and FMRP LCD (+ 7dG). Dot plots of the natural sequence rG4s by $log_2$ RBNS enrichment of **M)** G3BP1 (+ Guanine) and **N)** FMRP LCD (+ Guanine). RBNS enrichment for WT sequences are shown as large circles colored by natural rG4, specifically *VEGFA* (red), *U2AF2* (dark orange), *ARHGEF12* (orange), *EWSR1* (light orange), *HNRNPA2B1* (yellow), and *PABPN1* (grey) (Data is the average of 2 independent replicates). Mutated sequences for each rG4 are plotted as smaller circles behind the WT sequence, colored by the ΔWT scores determined above (red is destabilizing, positive, grey is zero, blue is stabilizing, negative).

### Supplemental Figure 6

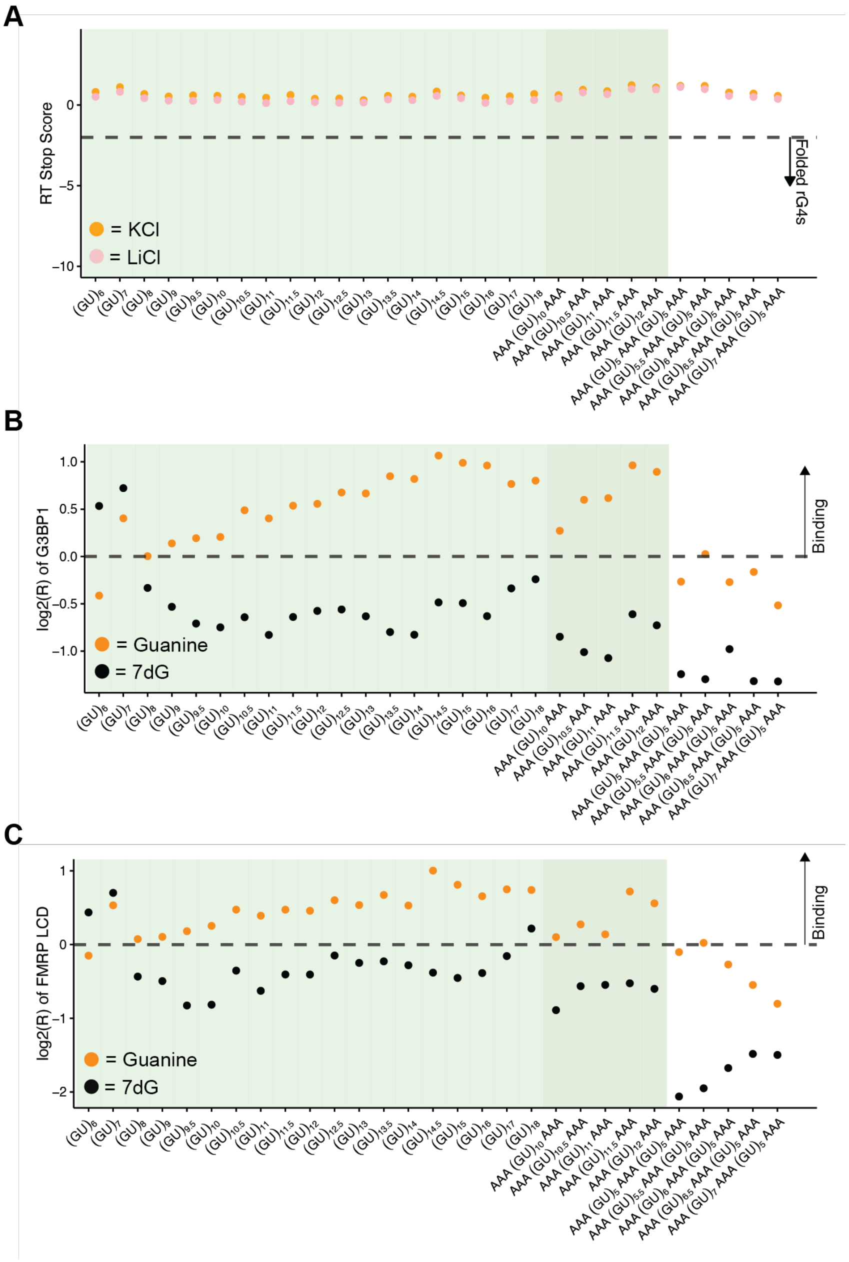


**Supplemental Figure 6. pUGs lack RT stops but bind proteins.** Dot plots comparing **A)** RT Stop Score in KCl (orange) or LiCl (pink), **B)** $\log_2(R)$ enrichment for G3BP1 (+ Guanine) (dark orange) or (+ 7dG) (black), and **C)** $\log_2(R)$ enrichment for FMRP LCD (+ Guanine) (dark orange) or (+ 7dG) (black). Light green rectangles correspond to putative pUG sequences varied by number of repeats, dark green to putative pUG sequences bookended by AAA sequences, and white for sequences not predicted to form pUGs. Dashed lines at -2 for RT Stop Score correspond to folded rG4 characterization, while lines at 0 for $\log_2(R)$ enrichment correspond to the threshold for improved binding/greater enrichment over input.
